# Distinct Virulence Mechanisms of *Burkholderia gladioli* in Onion Foliar and Bulb Scale Tissues

**DOI:** 10.1101/2024.10.07.617006

**Authors:** Sujan Paudel, Yaritza Franco, Mei Zhao, Bhabesh Dutta, Brian H. Kvitko

## Abstract

Slippery skin of onion caused by *Burkholderia gladioli* pv. *alliicola* (Bga) is a common bacterial disease reported from onion growing regions around the world. Despite the increasing attention in recent years, our understanding of the virulence mechanisms of this pathogen remains limited. In this study, we characterized the predicted genetic determinants of virulence in Bga strain 20GA0385 using reverse genetics approach. Using the closely related rice pathogen, *B. glumae* as a reference, comparative genomics analysis was performed to identify Bga candidate virulence factors and regulators. Marked and unmarked deletion mutants were generated using allelic exchange and the mutants were functionally validated using *in vitro* and *in vivo* assays. The role of mutants in pathogenic phenotypes was analyzed using onion foliar/seedling necrosis assays, the Red Scale Necrosis (RSN) assay and *in planta* bacterial population counts. The phytotoxin toxoflavin was a major contributor to foliar necrosis and bacterial populations whereas the type II and type III secretion system (T2SS/T3SS) were dispensable for foliar symptoms. In onion scale tissue, the T2SS single mutant *gspC* and its double and triple mutant derivatives all contributed to scale lesion area. Neither the lipocyclopeptide icosalide, toxoflavin, nor T3SS were required for scale symptoms. Our results suggest the quorum sensing *tofIMR* system in Bga regulates, toxoflavin, T2SS, and T3SS, contributing to onion symptom production. We show different virulence factors contribute to onion tissue-specific virulence patterns in Bga and that decreases in scale symptoms often do not result in decreased Bga populations in onion tissue.

## Introduction

Onion is one of the world’s most significant vegetable crops, cultivated on over 5 million hectares with an annual production of more than 100 million metric tons (Fayos et al., 2022). Bacterial diseases pose a substantial risk to the onion production as they can cause infection both in the field and during storage (Belo et al., 2023). Slippery skin of onion caused by *Burkholderia gladioli* pv. *allicola* (Bga) is one of the major diseases of onion in the United States (Schroeder et al., 2012). The pathogen is commonly isolated from onion fields and under storage conditions (Lee et al., 2005). Infected bulbs appear normal from the outside but are softer when pressed compared to the healthy ones. In the field, infection has been described to start from the neck of the onion and progress gradually to the basal plate before spreading to the neighboring bulbs (Wright et al., 1993). The rot caused by Bga is typically characterized by a watery, glassy appearance, whereas the rot associated with sour skin of onion, caused by the *Burkholderia cepacia* complex (Bcc), was described by Burkholder as slimy and yellow (Burkholder, 1950). Leaves infected by Bga developed water-soaked lesions with a bleached appearance, which often led to wilting and rot. Since its first discovery in the 1940s, the bacterium has been isolated and characterized from various onion growing regions in Asia, North and Central America, and Europe (Wright et al., 1993; Lee et al., 2005; Lamovšek et al., 2016; Félix-Gastélum et al., 2017). Despite its increasing economic significance to onion production, studies of the mechanisms of virulence in Bga are limited.

The comparatively well studied rice pathogen, *Burkholderia glumae* has been used as a useful reference model to infer virulence determinants in Bga because of their close phylogenetic relatedness (Vandamme et al., 2017). The ability of *B. glumae* to cause onion scale tissue necrosis is often utilized for genetic characterization studies (Karki et al., 2012). The well characterized virulence factors such as the phytotoxin toxoflavin, type II and type III secretion systems (T2SS/T3SS) are conserved in both *B. gladioli* and *B. glumae* (Fory et al., 2014; Seo et al., 2015; Lee et al., 2016). *In silico* analysis have also shown the presence of putative T3SS and T6SS in both *B. gladioli* and *B. glumae* (Fory et al, 2014; Seo et al., 2015). In *B. glumae,* the contribution of toxoflavin to rice chlorotic symptoms and wilting symptoms in various field crops has been demonstrated (Jeong et al., 2003; Seo et al., 2015). Similarly, T2SS and T3SS also contribute to rice panicles symptoms and bacterial populations in *B. glumae* (Goo et al., 2010; Karki et al., 2012). Likewise, the T2SS has also been shown to be an important virulence factor in *B. gladioli* pv. *agaricicola* (Chowdhury and Heinemann, 2006). The *tofIMR* quorum sensing system has been extensively studied as an important virulence regulator in both *B. glumae* and *B. gladioli* (Lee et al., 2016; Elshafie et al., 2019). The thiosulfinate tolerance gene (TTG) clusters, that contribute to enhanced allicin tolerance *in vitro*, also contribute to onion foliar symptoms and bacterial populations in Bga (Paudel et al., 2024b). Comparative genomics of *Burkholderia* strains isolated from various ecological niches revealed the broad distribution of the icosalide *icoS* nonribosomal peptide synthetase (NRPS). The *icoS* gene is present in some plant pathogenic *Burkholderia* species, including *B. cepacia*, *B. gladioli* pv. *agaricicola*, and *B. gladioli* pv. *gladioli* (Dose et al., 2018). Notably, this cluster is absent in rice-pathogenic *B. glumae* strains. In *B. gladioli* strain HKI0739, an endosymbiont, icosalide plays a protective role for its eukaryotic host against entomopathogens (Dose et al., 2018). Although the gene cluster is present in Bga, its potential role in onion host interactions remains unexplored.

All four *B. gladioli* pathovars, pv. *cocovenenans*, pv. *agariciola*, pv. *gladioli*, and pv. *alliicola* are capable of onion scale necrosis and the utility of the pathovar concept for *B. gladioli* has been questioned (Abachi et al., 2024). The *alliicola* pathovar, originally isolated from onion, forms a distinct evolutionary clade in *B. gladioli* species (Jones et al., 2021). Despite some sporadic studies on separate host systems, the genetic basis of virulence in Bga is only partially understood. In this study, we characterized the putative genetic determinants of virulence in onion isolated Bga strain 20GA0385 and empirically demonstrated that different virulence factors contribute to different symptoms in specific tissues, foliage vs. scale. Our results suggest that the phytotoxin toxoflavin is a critical virulence factor for onion leaf tissue whereas the T2SS was a significant contributor to scale lesion development. However, the reduction in Bga symptom production in scale tissue was not associated with corresponding decrease in bacterial population. The decoupling was evident with T2SS mutant *gspC* in scale tissue and icosalide mutant in leaf tissue.

## Results

### Putative virulence factors are conserved in *B. gladioli* pv. *alliicola* and *B. glumae*

In effort to characterize the candidate virulence factors in onion pathogenic Bga strain 20GA0385, a comparative genomics analysis was performed between Bga and well characterized *B. glumae* strain BGR1. The toxoflavin biosynthesis operon, T2SS, T3SS, and quorum sensing *tofIMR* system were all conserved in synteny between *B. glumae* BGR1 and Bga strain 20GA0385 (Supplementary Fig 1A-D). The toxoflavin biosynthesis polycistronic operon genes shared more than 90% identity between the two strains. The T2SS constituent genes also shared a high percentage identity ranging from 76-97%. The T3SS machinery genes were also highly conserved sharing percentage identity ranging from 70-97%. The Quorum Sensing (QS) genes also shared percentage identity from 80-97%. The antibiotic icosalide identified from a *B. gladioli* endosymbiont strain HKI0739 was absent from BGR1 but shared high percentage identity with a putative *icoS* gene found in 20GA0385 (Supplementary Table S1).

### Functional validation of mutants

Bacterial overnight suspensions were spotted onto a nitrocellulose membrane to enhance visual contrast of the yellow diffusible pigment toxoflavin when imaging the underside of LB agar plates. The toxoflavin biosynthesis mutant *toxA* lost yellow pigment production compared to the WT strain on LB media (Figure 1A). When the *toxA* open reading frame (ORF) was restored to the chromosome using allelic exchange, yellow pigment production was restored similar to that of the WT strain (Figure 1A).

**Figure 1:**
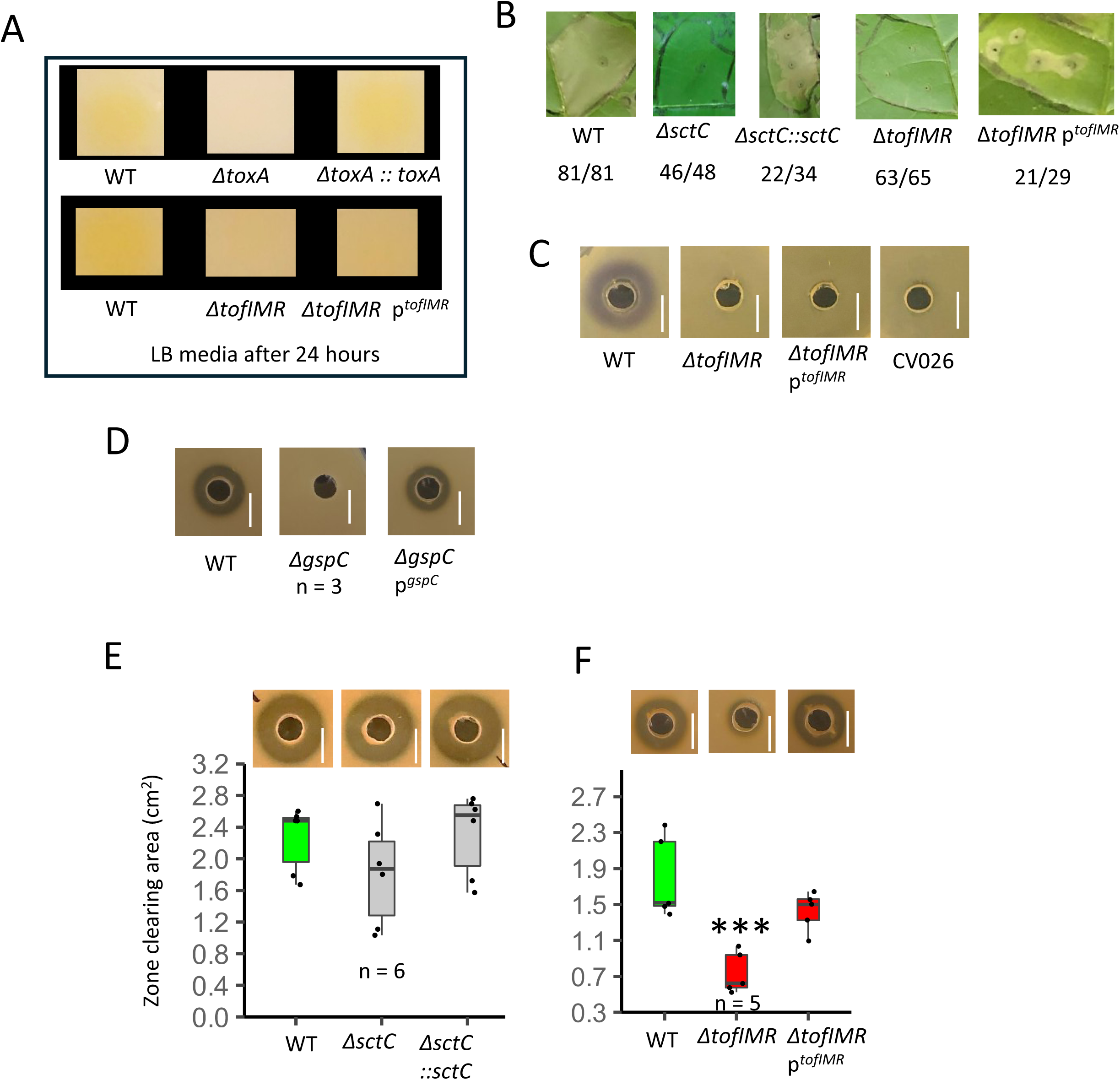
Functional validation of select mutants generated in the study. **A,** Visualization of diffusible yellow toxoflavin pigmentation from the back side of plate in the nitrocellulose membrane on top of LB agar plate spotted with **Top (**From left to right): WT, toxoflavin biosynthesis mutant *toxA* and *toxA* restoration clone; **Bottom** (from left to right: WT, quorum sensing mutant *tofIMR*, and *tofIMR* complementation clone. **B,** Tobacco cell death assay phenotype of different mutants compared to the WT strain. Representative image from one of the experimental repeats is shown. The ratio of infiltrated spots displaying the representative image phenotype to the total number of infiltrated spots is shown at the bottom of each strain.. **C,** *Chromobacterium* AHL reporter assay showing the purple violacein phenotype of (left to right) WT, Δ*tofIMR*, *tofIMR* complementation clone and CV026 negative control. **D,** Casein protease plate assay showing the clearing zone phenotype of tested (from left to right) WT, T2SS mutant *gspC* and the *gspC* plasmid complementation clone. n = total number of observations. Representative image from experimental repeats is shown for all the *in vitro*/*in vivo* assays. Box plot image with quantification of zone clearing area from the protease plate assay for **E,** WT, T3SS mutant *sctC* and *sctC* restoration clone and **F,** WT, QS mutant *tofIMR* and *tofIMR* complementation clone. n represents a total of observation. Representative image from experimental repeats presented at the top of corresponding box plot. The test of significance was performed using pairwise t-test function in RStudio. Level of significance: 0 ‘***’ 0.001.

The functional validation of T3SS was done using a tobacco cell death assay. The tobacco cell death area of T3SS Δ*sctC* mutant was visually compared to the WT strain. The cell death area of the *sctC* mutant was consistently reduced compared to the WT strain. The chromosomally restored *sctC* clone complemented the cell death phenotype of the *sctC* mutant (Figure 1B). The T2SS Δ*gspC* had comparable tobacco cell death phenotypes to the WT strain. The Δ*gspC*Δ*sctC* double and Δ*toxA*Δ*gspC*Δ*sctC* triple mutant strains were reduced in cell death area compared to the WT strain (Supplementary Figure S2E).

The *Chromobacterium* Acyl Homoserine Lactones (AHL) assay was performed to functionally validate the *tofIMR* mutant *in vivo* using CV026 reporter strain (McClean et al., 1997). The AHL molecules produced by the test treatment when recognized by the CV026 reporter strain produce violacein ring around the area of inoculation. The normalized suspension of the WT strain when inoculated in the CV026 lawn produced violet ring around the zone of inoculation whereas the *tofIMR mutant* did not (Figure 1C). Surprisingly, the plasmid based *tofIMR* complementation clone did not restore violacein production in the CV026 lawn background although other *tofIMR* phenotypes including *in planta* phenotypes were complemented as described below (Figure 1C).

The T2SS mutant strain was validated using Casein hydrolysis protease plate assay. The WT strain produced a clearing area around the hole of inoculation on dry milk augmented agar plates whereas the T2SS Δ*gspC* lost the clearing phenotype (Figure 1D). The plasmid pBBR1MCS-2 with *gspC* ORF when expressed in *gspC* mutant background partially restored the clearing area suggesting the T2SS secreted proteases cause hydrolysis of casein in the agar plate (Figure 1D). The Δ*gspC*Δ*sctC*, Δ*gspC*Δ*icoS*::Gm^R^*gus*, and Δ*toxA*Δ*gspC*Δ*sctC* double and triple mutant strains retained the *gspC* single mutant clearing zone phenotype (Supplementary Figure S2B). The clearing area for the T3SS *sctC* mutant was variable across the experimental repeats. Across a total of six observations, the clearing area for the *sctC* mutant strain was not significantly different from the WT strain (Figure 1E). Clearing area quantification was not performed for the *gspC* single mutant or its double and triple mutant derivatives as a consistent visual phenotype was observed across all experimental repeats.

The swarming motility assay was done to visually compare the *in vitro* swarming behavior of icosalide single mutants to the WT strain. Both the triple stop codon interruption mutant Ψ*icoS* and the Δ*icoS*::Gm^R^*gus* were reduced in swarming area compared to the WT strain (Supplementary Figure S2F). Surprisingly, the Δ*gspC*Δ*icoS*:: Gm^R^*gus* double mutant was further reduced in swarming area compared to the icosalide single mutants. The T2SS *gspC* mutant produced a bigger swarming area compared to the two icosalide single mutants (Supplementary figure S2F).

### The regulation of toxoflavin production by QS *tofIMR* system is dependent upon media conditions

We had observed variation in yellow pigmentation based on growth media. To test if the Bga 20GA0385 *tofIMR* system behaves similarly, the toxoflavin production by *tofIMR* mutant was visualized and compared to WT strain on LB, LM, and MGY media agar. On LB agar, diffusible yellow pigmentation was not observed in *tofIMR* spotted membranes in 4 out of 5 repeats (Figure 1A). On the MGY media, the *tofIMR* spotted membrane appeared yellow like the WT spotted membrane (Supplementary figure S2A). The variation in *tofIMR* pigmentation was more pronounced on LM media across the repeats. For two experimental repeats in LM media, the differentiation in yellow pigmentation between the WT and *tofIMR* mutant spotted membrane was not visually distinct 24 hours post inoculation (Supplementary Figure S2B). After an additional 36 hours of incubation at room temperature, the differences in yellow pigmentation between the treatments became more distinct (Supplementary Figure 2C). The restoration of yellow pigment in the *tofIMR* complementation clone was more prominent on the LM agar plate compared to the LB plate. (Supplementary Figure S2B; Figure 1A).

### The QS *tofIMR* system regulates protease secretion, tobacco cell death and swarming motility in Bga strain 20GA0385

The QS *tofIMR* mutant was also tested in the casein protease plate assay and tobacco cell death assay to assess its role in regulation of T2SS- and T3SS-associated phenotypes, respectively. The mutant was significantly reduced in casein hydrolysis protease plate clearing area compared to the WT strain. The *tofIMR* complementation clone partially restored the *tofIMR* casein milk agar clearing area to the WT level (Figure 1F). The *tofIMR* inoculated tobacco leaves were reduced in cell death area compared to the WT strain (Figure 1F). The *tofIMR* complementation clone partially restored the cell death area of *tofIMR* mutant (Figure 1B).

In the swarming motility assay, the QS *tofIMR* mutant was restricted around the point of inoculation on the plate whereas the WT strain swarmed to the edge of the plate (Supplementary Figure S2F). The *tofIMR* complementation clone had a dramatically increased swarming area compared to the *tofIMR* mutant (Supplementary Figure S2F).

### The phytotoxin toxoflavin is a key virulence factor in onion leaf tissue

The phytotoxin toxoflavin biosynthesis mutant *toxA* was significantly reduced in onion necrosis length and bacterial population compared to the WT (Figure 2A, B). In a previous study, we reported the positive contribution of the thiosulfinate tolerance gene clusters (TTG) to onion foliar necrosis and bacterial populations in Bga strain 20GA0385 (Paudel et al., 2024). The *toxA* mutant in the TTG mutant background was also significantly reduced in both foliar lesion length and bacterial populations compared to the WT strain (Figure 2C, D). The reduction in necrosis length and bacterial population of *toxA* mutant was restored to WT levels with *toxA* chromosomal restoration clone (Supplementary Figure S4A, B). The icosalide *icoS* triple stop codon interruption mutant Ψ*icoS* and Gm^R^*gus* marked deletion mutants were both unchanged in foliar necrosis length compared to the WT (Figure 2E, G). The two mutants, however, were significantly reduced in leaf bacterial populations *in planta* compared to the WT (Figure 2F, H). In the onion scale tissue, the Δ*toxA* and Δ*toxA*ΔTTG double mutants were not altered in necrosis area and bacterial populations compared to the WT strain (Figure 2I, J; Supplementary Figure 3A, D). Similarly, the icosalide single mutants, Ψ*icoS* and Δ*icoS::*Gm^R^*gus* were both unchanged in RSN area and bacterial populations (Figure 2K, L; Supplementary Figure 3B, C).

**Figure 2:**
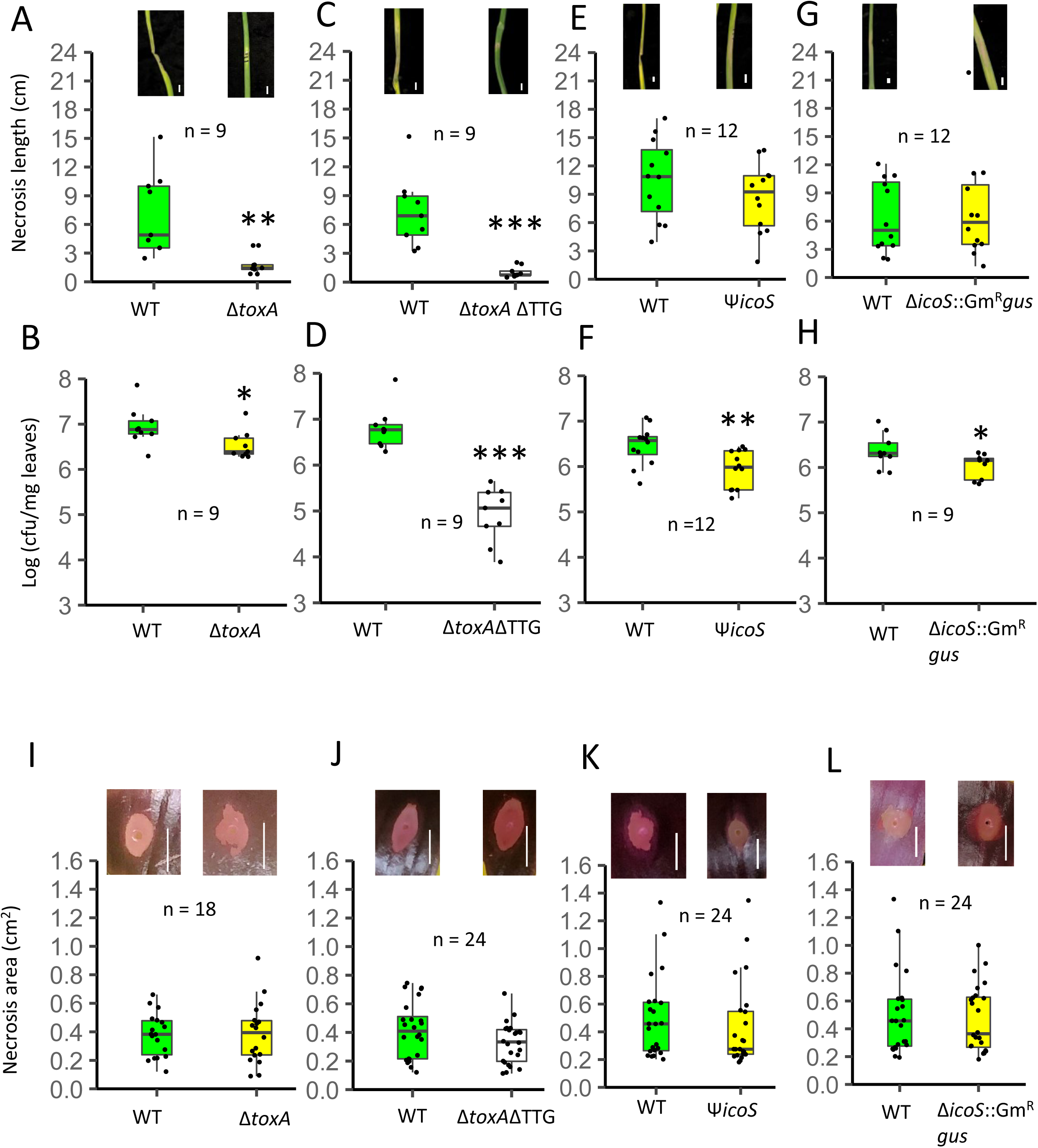
The phytotoxin toxoflavin in Bga strain 20GA0385 contributes to foliar necrosis and bacterial populations. Box plot showing onion foliar necrosis length and *in planta* bacterial populations of Bga WT 20GA0385 strain compared to **A**, **B,** *toxA*, **C, D,** Δ*toxA*ΔTTG **E, F,** Ψ*icoS* **G, H,** Δ*icoS*::Gm^R^*gus* mutants. Box plot showing RSN area of Bga WT 20GA0385 strain compared to **I** Δ*toxA*, **J,** Δ*toxA*ΔTTG, **K,** Ψ*icoS*, **L,** Δ*icoS*::Gm^R^*gus* mutants. A representative image of infected leaves or scales inoculated with treatments at 3 dpi is presented above the box plot. n is the total number of observations across the experimental repeats. The test of significance between the necrosis length/area and bacterial populations was performed using pairwise t-test function in RStudio. Level of significance: 0 ‘***’ 0.001 ‘**’ 0.01 ‘*’ 0.05. Scale: 1 cm.

### The T2SS contributes to onion scale necrosis

The T2SS mutant *gspC* and T3SS Δ*sctC* were tested for their virulence role using the foliar and RSN phenotypic assays. The *gspC* and *sctC* single mutants both produced comparable leaf lesion length and *in planta* populations compared to the WT strain (Figure 3A, B; Supplementary Figure S4C, D). In the RSN assay, the *gspC* mutant was reduced in necrosis area while the *sctC* mutant was unchanged relative to the WT strain (Figure 3D, E). The *gspC* sequence region when expressed in the *gspC* mutant background using the expression plasmid pBBR1MCS-2 partially restored the phenotype (Figure 3D). The Δ*gspC*Δ*sctC* and Δ*gspC*Δ*icoS*::Gm^R^*gus* double mutants both resembled the *gspC* single mutant in RSN phenotypes and foliar necrosis symptoms (Figure 3F, 3G, 3C, 3H). The *gspC*/*sctC* double mutant was also not impaired in leaf bacterial populations compared to the WT (Supplementary Figure S4E). The Δ*gspC*Δ*icoS*::Gm^R^*gus* population recovery from inoculated leaves was similar to the Δ*icoS*::Gm^R^*gus* single mutant (Figure 3I). The Δ*toxA*Δ*gspC*Δ*sctC* triple mutant when inoculated in onion seedling produced significantly shorter lesions length compared to the WT strain (Figure 4A). The mutant was also significantly reduced in seedling foliar population recovery compared to the WT (Figure 4B). Unexpectedly, the Δ*toxA*Δ*gspC*Δ*sctC* triple mutant did not recapitulate the red scale phenotype of the *gspC* single mutant (Figure 4C). To investigate whether this discrepancy was due to high initial bacterial inoculum concentration (OD_600_ = 0.7), we conducted the same test using 100-fold diluted inoculum. With the diluted bacterial suspension, the *gspC* mutant, and all *gspC* derivative strain were significantly reduced in RSN area compared to the WT strain (Figure 5). All the tested protein secretion system mutants and their double and triple mutant derivatives were unchanged in red onion scale bacterial populations relative to the WT strain at both high and low bacterial inoculum concentrations (Supplementary Figure 3E-I, 3K-N).

**Figure 3:**
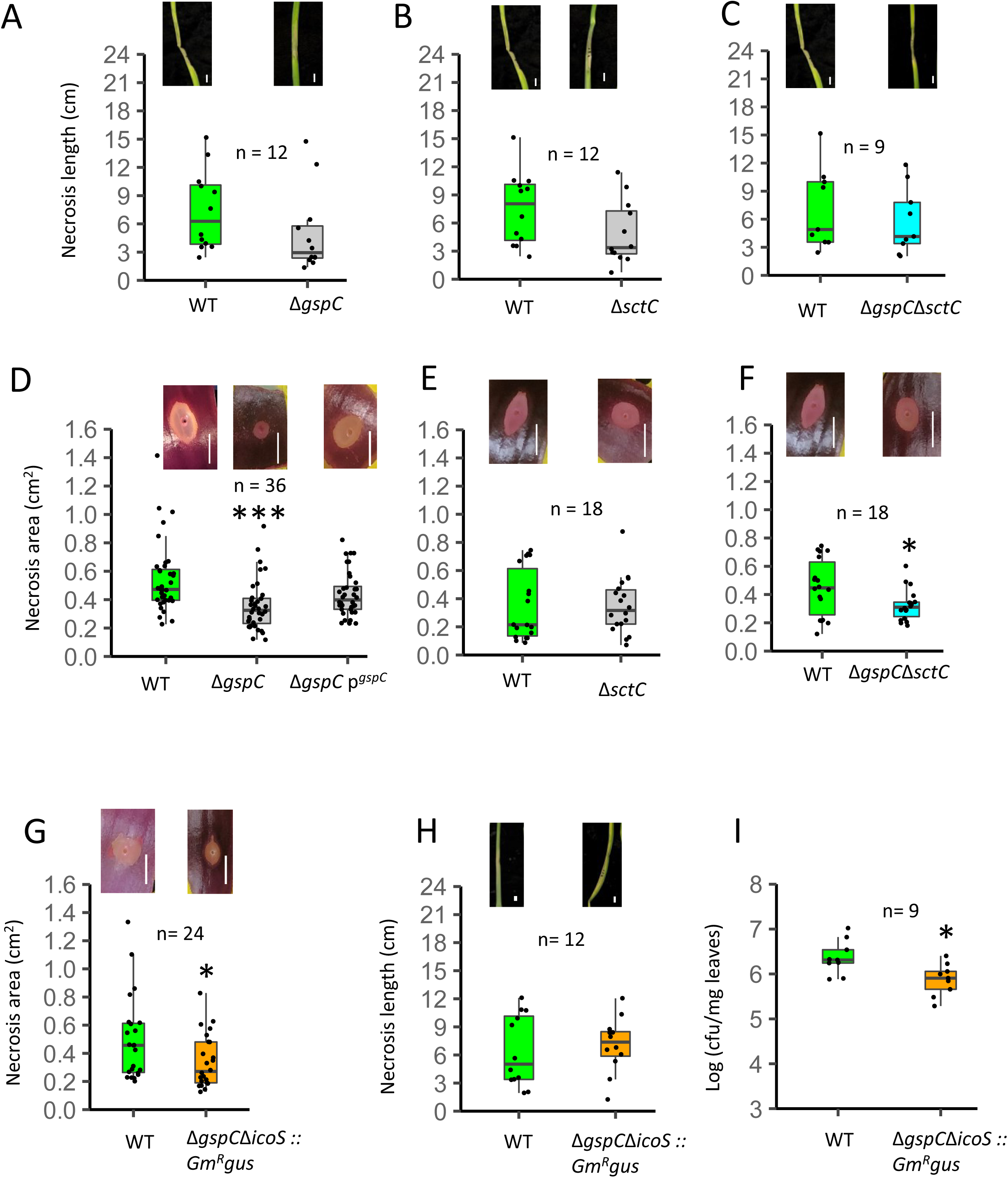
The T2SS contributes to RSN area but is not required for foliar symptoms. Box plot showing onion foliar necrosis length of Bga WT 20GA0385 strain compared to **A,** T2SS Δ*gspC*, **B,** T3SS Δ*sctC*, and **C,** Δ*gspC*Δ*sctC* mutants. Box plot showing RSN area of Bga WT 20GA0385 strain compared to **D,** T2SS Δ*gspC* and *gspC* plasmid complementation clone, **E,** T3SS Δ*sctC*, **F,** Δ*gspC* Δ*sctC* and **G,** Δ*gspC*Δ*icoS*::Gm^R^*gus* mutants. Box plot showing the comparison of **H,** necrosis length and **I,** leaf bacterial populations between WT and Δ*gspC* Δ*icoS*::Gm^R^*gus* mutant. A representative image of infected leaves or scales inoculated with treatments at 3 dpi is presented above the box plot. n is the total number of observations across the experimental repeats. The test of significance between the necrosis length/area and bacterial populations was performed using pairwise t-test function in RStudio. Level of significance: 0 ‘***’ 0.001 ‘**’ 0.01 ‘*’ 0.05. Scale: 1 cm.

**Figure 4:**
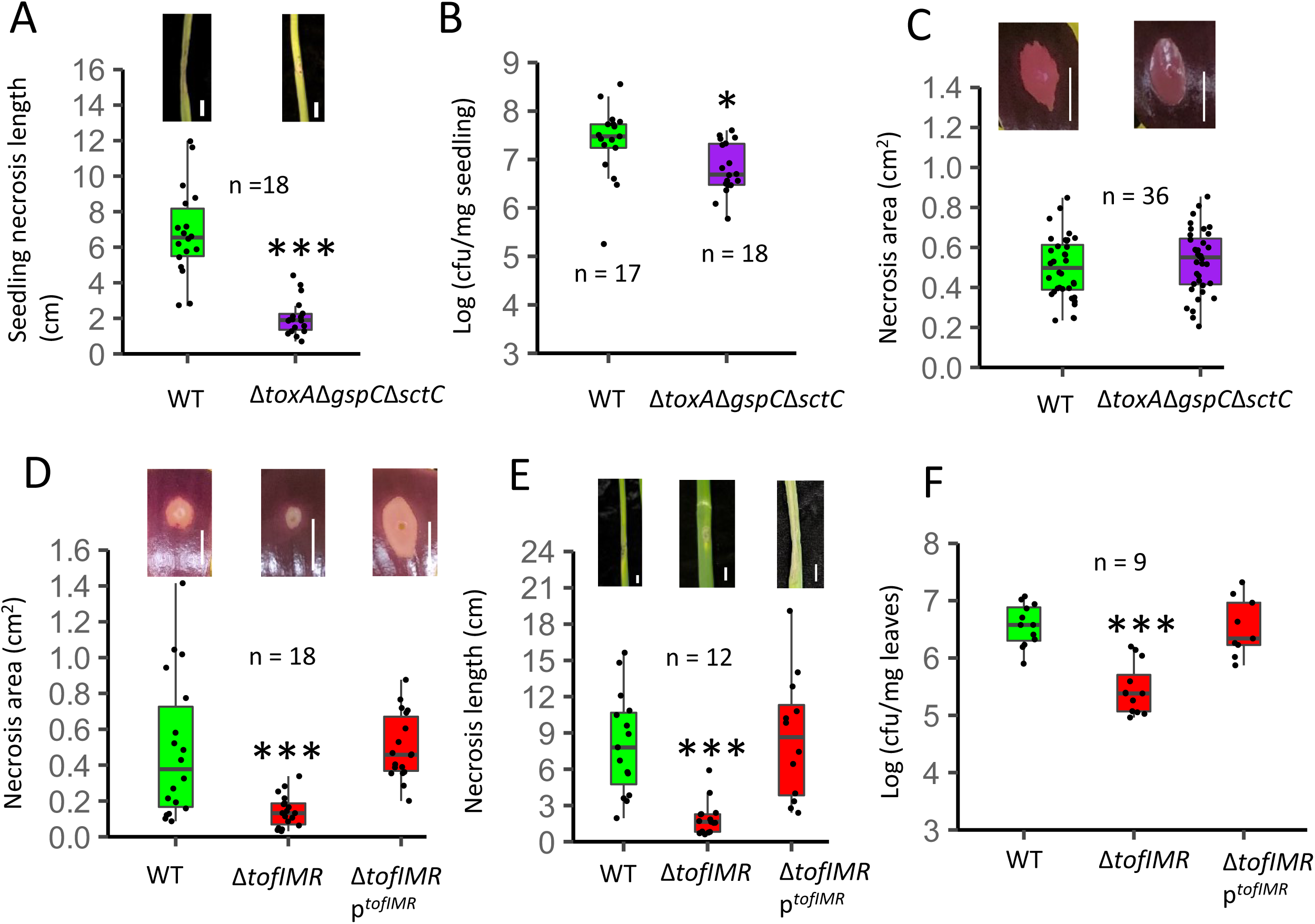
Box plot showing the comparison of **A,** seedling lesion length, **B,** seedling bacterial populations and **C,** RSN area between the WT and the Δ*toxA*Δ*gspC*Δ*sctC* triple mutant derivative. A representative image of inoculated seedling/scale at 3 dpi is shown above the box plot. **D-F**, **The *tofIMR* QS system regulates virulence phenotype of Bga strain 20GA0385**.Box plot showing the comparison of **D,** RSN area, **E,** foliar necrosis length, and **F,** foliar in planta populations between the WT, QS mutant *tofIMR* and the *tofIMR* plasmid complementation clone. n is the total number of inoculated samples across the experimental repeats. The test of significance between the necrosis length/area and bacterial populations was performed using pairwise t-test function in Rstudio. Level of significance: 0 ‘***’ 0.001 ‘**’ 0.01 ‘*’ 0.05. Scale: 1 cm.

**Figure 5:**
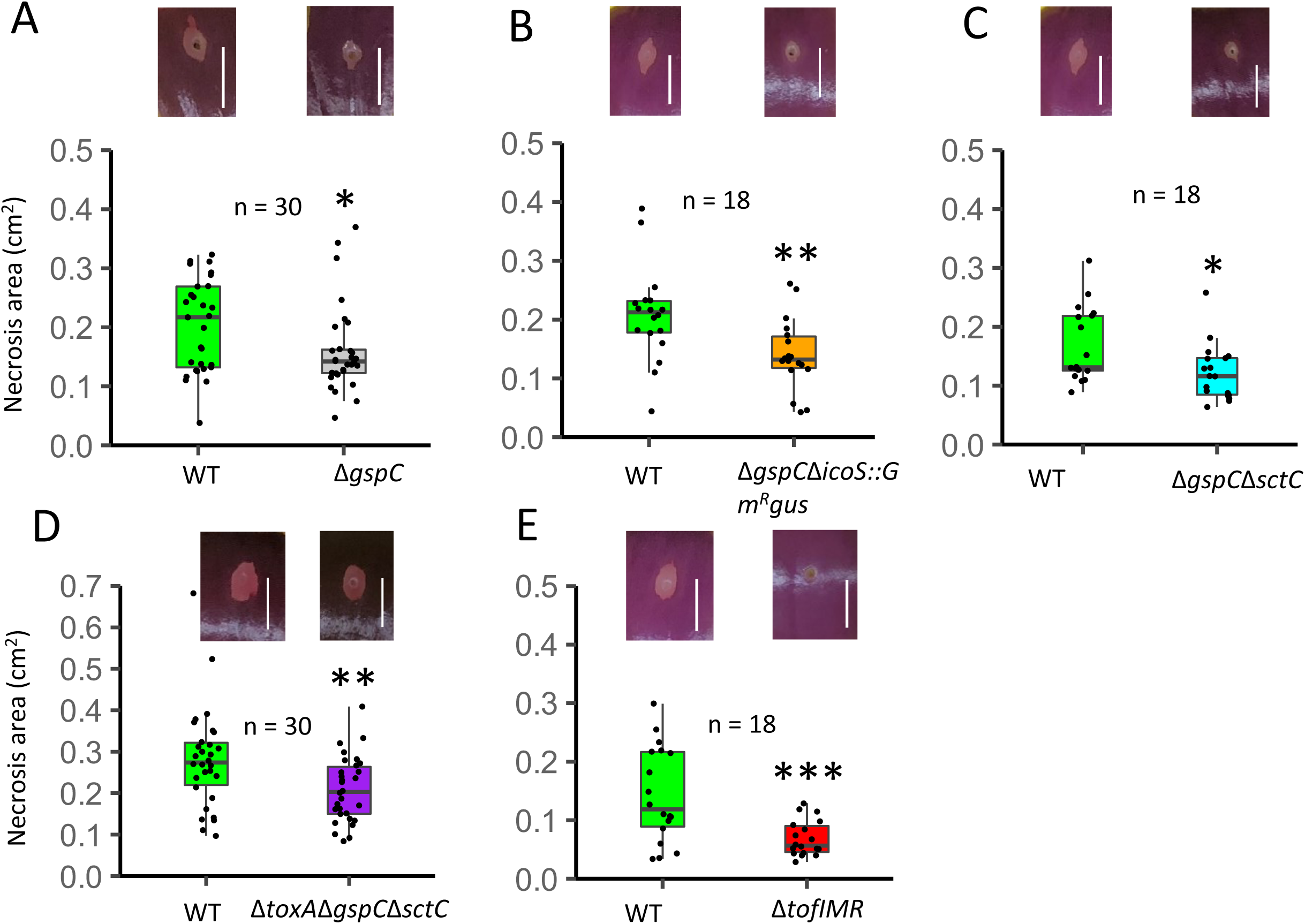
The virulence role of T2SS in onion scale is maintained in 100-fold diluted starting bacterial concentration. Box plot showing the RSN area of WT strain at 3 dpi compared to **A,** Δ*gspC*, **B,** Δ*gspC* ΔicoS::Gm^R^*gus*, **C,** Δ*gspC*Δ*sctC*, **D,** Δ*toxA*Δ*gspC*Δ*sctC***, E**, Δ*tofIMR* mutants. A representative image of infected leaves or scales inoculated with treatments at 3 dpi is presented above the box plot. n is the total number of observations across the experimental repeats. The test of significance between the necrosis length/area and bacterial populations was performed using pairwise t-test function in RStudio. Level of significance: 0 ‘***’ 0.001 ‘**’ 0.01 ‘*’ 0.05. Scale: 1 cm.

### The QS *tofIMR* system regulates onion symptoms production

The QS *tofIMR* system that regulates many physiological and virulence traits in *Burkholderia* species was also tested for its contribution to virulence and bacterial population phenotype in RSN and foliar assays. The *tofIMR* mutant significantly contributed to RSN area (Figure 4D). The *tofIMR* mutant when expressed with *tofIMR* plasmid complementation clone restored the RSN area to the WT levels (Figure 4D). Similarly, the *tofIMR* mutant was also significantly reduced in foliar necrosis length and bacterial populations compared to the WT. The *tofIMR* complementation clone in pBBR1MCS-2 plasmid restored the RSN area and bacterial populations phenotype to the WT level (Figure 4E, F).

## Discussion

We characterized the contributions of putative virulence determinants to Bga onion leaf and scales symptoms production and in planta bacterial populations. Toxoflavin make major contributions to foliar symptoms production and bacterial populations in leaves. While the T3SS had little effect on Bga virulence, the T2SS was necessary for symptom development in onion scales but did not contribute to foliar necrosis or bacterial population growth. Results from both foliar and RSN phenotypic assays suggest that the *tofIMR* quorum sensing system is a key virulence regulator in Bga strain 20GA0385 (Figure 4D-F). The *in vitro* and *in vivo* experimental results from protease and tobacco cell death assays are consistent with *tofIMR* regulation of T2SS and T3SS in Bga strain 20GA0385 (Figure 1A-B, F). In the phenotypic assays, we routinely observed decoupling of bacterial populations and onion tissue symptoms. This trend was more pronounced in onion scale tissue as compared to foliar tissue (Figure 3D-G; Figure 5; Supplementary Figure S3). Our study suggests tissue specific virulence factors in Bga onion virulence.

The phytotoxin toxoflavin is an extensively studied virulence factor in *B. glumae.* The toxin contributes to foliar chlorosis and necrotic symptoms in rice leaf tissue (Ilyama et al., 1995; Suzuki et al., 1998; Yoneyama et al., 1998). The studies pertaining to the role of toxin in Bga onion scale and leaf symptoms production are limited. In our study, we found the toxin is a major contributor to Bga onion foliar symptoms (Figure 2A). In the Bga WT strain inoculated onion leaves, the symptoms appeared as a greyish lesion after 24-36 hours near the point of inoculation and gradually spread away from the point of inoculation. After 3-4 days, the lesion extended towards the tip of the blades and the leaf appeared wilted. The *toxA* mutant inoculated leaf samples in general produced lesions limited around the point of inoculation and rarely progressing towards the blade tip. The *toxA* mutant was also significantly reduced in leaf bacterial population *in planta* compared to WT (Figure 2B). However, the reduction in population was not dramatic. In scale tissue, the necrosis symptoms started appearing from the point of inoculation and developed radially away from the wounding site typically appearing as an oval shaped lesion 3-4 days post inoculation (dpi). Despite being reported as a major virulence factor in rice *Burkholderia* system by multiple independent studies, the lack of clear toxoflavin contribution to onion scale symptoms was surprising (Figure 2I).

TTG clusters have been characterized in several onion bacterial pathogens including *P. ananatis*, as well as *B. cepacia*, *B. orbicola*, and *B. gladioli* (Stice et al., 2020; Paudel et al., 2024b). The clusters in these species confers *in vitro* tolerance to allicin, a volatile antimicrobial defensive organosulfur compound produced by garlic. In a previous study, we demonstrated the contribution of TTG clusters in Bga strain 20GA0385 to onion foliar leaf symptoms and bacterial populations (Paudel et al., 2024b). While the *toxA* single mutant was slightly reduced in *in planta* foliar population recovery, the Δ*toxA*ΔTTG double mutant population recovery from leaf tissue was further reduced compared to *toxA* single mutant (Figure 2B, D). The foliar population recovery of the Δ*toxA*ΔTTG double mutant was similar to that of the TTG single mutant reported previously (Paudel et al., 2024b). The *toxA* mutant was also severely reduced in leaves necrosis compared to the WT (Figure 2A). Despite the dramatic reduction, the *toxA* mutant still produced a necrotic lesion mostly centered around the point of inoculation. As the TTG cluster single mutant was also severely reduced in leaf bacterial population (Paudel et al., 2024b), toxoflavin independent necrosis factor could also be present in Bga strain. By contrast, both *toxA* and TTG cluster single mutants and the *toxA*/TTG double mutant did not contribute to RSN area nor populations.

The lipocyclopeptide antibiotic icosalide was described in the beetle endosymbiont *B. gladioli* strain HKI0739. The non ribosomal peptide synthetase (NRPS) assembled secondary metabolite protected the beetle against entomopathogenic bacteria and inhibited swarming motility of the HKI0739 (Dose et al., 2018). A putative icosalide homolog *icoS* was also found in the Bga strain 20GA0385 that shared a high identity percentage with the biosynthetic NRPS gene described previously (Supplementary Table S4). As the ORF encoding the putative icosalide biosynthetic gene is over 15 kb size, full gene complementation was not pragmatic. Thus, two independent mutations were generated to assess the virulence role of icosalide in 20GA0385. Both mutants were unchanged in their foliar and onion scale necrosis and onion scale populations (Figure 2E, G, K, L). The two mutants did contribute significantly to foliar populations (Figure 2F, H). As the T2SS *gspC* mutant contributed to RSN area, a double mutant was created by introducing *gspC* mutant into the Δ*icoS*::Gm^R^*gus* background to check if it has further reduction in the RSN phenotype. The double mutant was also significantly reduced in RSN symptoms as the *gspC* single mutant (Figure 3G). The role of Bga icosalide in swarming motility of the bacterium was analyzed by developing an *in vitro* swarming assay. In contrast to what was observed with the strain HKI0739 where mutants displayed increased swarming, both icosalide mutants were visually reduced in swarming area compared to the WT strain. A further reduced swarming area was observed for the *gspC/icoS*::Gm^R^*gus* double mutant compared to both single mutants (Supplementary Figure S2F). While the icosalide mutants in HKI0739 demonstrate increased swarming behavior, our results indicate the opposite phenotype in Bga strain 20GA0385. Despite sharing high homology, the strain specific differences observed in the *in vitro* swarming phenotype between the two strains was noteworthy. The differences in ecological niche of the two strains harboring the icosalide cluster might have caused the differences in swarming behavior of the strains. Foliar assay results suggest the Bga icosalide cluster mutants recovery from infected leaf tissue was significantly lower compared to the WT strain (Figure 2F, H). As authors demonstrated the antibiotic activity of icosalide in HKI0739 strain suggesting its potential role in eukaryotic host protection, the Bga icosalide cluster could also help the bacteria interact with competitors promoting the colonization of bacteria in onion leaf tissue. Further experimental tests are needed to test this hypothesis.

The virulence-associated protein secretion systems, particularly T2SS, and T3SS have also been characterized as a virulence factor in *B. glumae* and *B. gladioli.* T2SS in *B. glumae* contributes to rice panicles symptoms and *in planta* populations and in *B. gladioli*, T2SS is required for mushroom tissue necrosis (Chowdhury and Heinemann, 2006; Devescovi et al., 2007; Kang et al., 2008; Goo et al., 2010). The role of *B. gladioli* T3SS in mycophagy has also been demonstrated (Swain et al., 2017). We found that neither T2SS nor T3SS contributed to foliar necrosis nor *in planta* bacterial populations (Figure 3A, B; Supplementary Figure S4C, D). The T2SS mutant *gspC* and its double mutant derivatives all contributed positively to red scale necrosis symptoms production in both high (∼6 x 10^6^ CFU) and low (∼6 x 10^4^ CFU) inoculum (Figure 3D, E; Figure 5A-D). Surprisingly, the Δ*toxA*Δ*gspC*Δ*sctC* triple mutant showed a similar RSN area to the WT strain when higher initial bacterial concentration was used (Figure 4C). All the mutants that contributed to RSN area at higher inoculum concentration also showed similar results at lower inoculum concentration (Figure 5A-D). Consistent with the findings from *B. glumae* and *B. gladioli* pv. *agaricicola* studies, the T2SS in Bga also functions as an important virulence factor in onion scale tissues.

The decoupling of Bga *in planta* bacterial populations and tissue symptoms was a common trend observed in the study. In leaf tissue, the secondary metabolite icosalide was not required for leaf necrosis but contributed to bacterial populations (Figure 2E-H). In the scale tissue, none of the tested mutants that contributed to necrosis area had reduced *in planta* bacterial populations (Supplementary Figure S3). Additional repeats were done for mutants that were reduced in onion scale necrosis with 100-fold diluted starting concentration. All tested mutants at lower concentration were still recovered at similar level to the WT strain despite retaining the RSN area phenotype at higher concentrations (Supplementary Figure S3 K-O; Figure 5A-D).

The quorum sensing *tofIMR* system has been shown to regulate multiple virulence factors in *B. glumae* (Devescovi et al., 2007; Goo et al., 2010; Lelis et al., 2019). Upon visual examination across three different media conditions, variation in toxoflavin pigmentation between the WT and *tofIMR* mutant was observed across the experimental repeats. The variation was more pronounced in LM media compared to LB and MGY media (Supplementary Figure S1B, C). The visual complementation of yellow pigmentation by *tofIMR* plasmid-based complementation clone was more pronounced on LM media compared to LB (Supplementary Figure S1B, C; Figure 1A). The regulators that contribute to *tofIMR* independent production of toxoflavin have been described previously (Chen et al., 2015). As described previously, we also observed variation in pigmentation of toxoflavin based on inoculation methods and media conditions (Chen et al., 2012). Through the *in vitro* assays we also demonstrated that the *tofIMR* system regulates the function of T2SS, T3SS, and swarming motility in Bga (Figure 1 B, F; Supplementary. Figure S3A-C, F). The virulence phenotype of *tofIMR* mutant in both leaves and scale was complemented by *tofIMR* complementation clone (Figure 4D-F). Our results from multiple independent experiments suggest that quorum sensing *tofIMR* system regulates multiple virulence traits in Bga strain 20GA0385.

Among the tested virulence factors, only T2SS *gspC* contributed to red scale symptoms production. The *gspC* mutant still produced noticeable red scale necrosis symptoms relative to the *tofIMR* system suggesting other unknown factors could yet contribute to RSN phenotype. The *tofIMR* system may regulate additional unknown virulence factors.

In summary, we observed the phytotoxin toxoflavin and the T2SS play onion tissue-specific virulence roles in Bga strain 20GA0385. Despite the key role of toxoflavin in leaf tissue necrosis, its lack of a clear role in red scale necrosis was unexpected. Likewise, commonly studied protein secretion systems T2SS and T3SS were not required for foliar symptoms production or bacterial population growth. No combined effect or synergism of individual virulence factors on symptom production or bacterial population was observed with Bga in our study. Scale assay necrosis area results suggest additional unknown virulence factors/regulators could contribute to onion scale symptoms. Furthermore, the decoupling of bacterial populations vs necrosis area particularly in the onion scale tissue suggests the bacterium is a well-adapted scale/bulb colonizer utilizing multiple virulence factors to infect the onion host. Future studies should focus on identification of additional virulence factors that contribute to onion symptom production and bacterial population in bulb tissues. Different management strategies would be required for the control of Bga foliar versus bulb symptoms.

## Materials and Methods

### Bacterial growth conditions

*Escherichia coli* strains and plasmids used for allelic exchange and construction of the complementing strains are listed in Supplementary Table S2. Primers and synthesized dsDNA fragments used in the study are listed in Supplementary Table S3. The mutants and the complementation clones generated in this study are listed in Supplementary Table S4. *Escherichia coli* strains DH5α, Mah1, RHO5, and Bga strain 20GA0385 used to generate mutants were grown in LB (per liter, 10 g of tryptone, 5 g of yeast extract, 5 g of NaCl) broth or LM (per liter, 10 g tryptone, 6 g yeast extract, 1.193 g KH_2_PO_4_, 0.6 g NaCl, 0.4 g MgSO_4_.7H_2_0) at 37°C and 28°C, respectively. Liquid broth was supplemented with Difco Agar (per liter, 15 g for LB and 18 g for LM) to make solid media. Antibiotics and chemicals were supplemented with the growth media at the following final concentrations, per milliliter: 50 µg of kanamycin (Km), 10 µg of gentamicin (Gm), 100 – 200 µg of Diaminopimelic acid (DAP), 50 µg of X-Gal (5-bromo-4-chloro-3-indolyl-beta-D-galacto-pyranoside), 25 µg of chloramphenicol (Cam), 100 µg of Xgluc (5-bromo-4-chloro-3-indolyl-beta-D-glucuronic acid), and 40-60 µg of rifampicin, as appropriate.

### Plant growth conditions

Onion sets (*Allium cepa L.* cv. Century) were planted in 10 cm x 8 cm (diameter x height) plastic pots filled with SunGrow 3B potting soil and maintained at greenhouse conditions with 25–28^0^C, 12L:12D photoperiod for 5 months from January to May until inoculation. To grow the seedlings, onion seeds (*Allium cepa* var. Texas Grano 1015Y) were sown in 3-inch pots with Sungrow 3B potting soil and were maintained in growth chamber conditions at ∼80^0^F, 40-50% humidity and watered every 2-3 days.

### Recombinant DNA techniques

PCR was conducted using GoTaq Green polymerase (Promega, Fitchburg, WI) or Q5 High-Fidelity DNA polymerase (New England Biolabs, Ipswich, MA) following manufacturer’s protocols. Primers and double stranded DNA were ordered/synthesized from Eurofins Genomics LLC (Louisville,KY) and Twist Biosciences (South San Franscisco, CA) respectively. Plasmid DNA was purified using GeneJet Plasmid Miniprep Kit (Thermoscientific, Watham, WA). PCR product and Gel cleanup was performed using Monarch NEB PCR cleanup kit and Monarch NEB Gel extraction kit (New England Biolabs), respectively. Restriction digest was done with enzymes purchased from New England Biolabs following manufacturer’s recommendations. Gateway recombination was done with Gateway BP or LR Clonase II enzyme mix from Thermo Fisher Scientific following manufacturer’s recommendations. Gibson cloning was performed with Gibson Assembly Master Mix (2X) from New England Biolabs. Gateway reaction products were transformed into chemically competent DH5α cells unless noted otherwise. The purified plasmid with the deletion construct was then transformed into electro-competent RHO5 cells. Sanger sequencing of PCR product and plasmids was done with Eurofins Genomics LLC (Louisville, KY). Whole-plasmid sequencing was done at Plasmidsaurus (Plasmidsaurus, Eugene, OR). Sequences were analyzed using Geneious Prime v. 2023.2.1.

### Identification of putative virulence factors in Bga strain 20GA0385

The rice pathogenic strain BGR1 (GenBank: GCA_000022645.2) was used as a reference to identify putative virulence factors and regulators in Bga strain 20GA0385 (Genebank: GCA_034365665.1). Another Bga strain FDAARGOS_389 was also included in the analysis as it had a closed genome available in the NCBI GeneBank database. The chromosome and plasmid sequences for each strain were downloaded from NCBI GenBank database and imported into the Geneious Prime v.2023.2.1 software. Sequences for *B. glumae* virulence factors were identified from the literature and map to reference plugin and/or custom BLAST feature was used in Geneious Prime v.2023.2.1 to locate corresponding sequence regions in FDAARGOS_389 and Bga strain 20GA0385. Obtained sequence information was utilized for designing the deletion construct and other downstream processes.

The gene synteny diagram and amino acid identity percentage was calculated using the Clinker CAGECAT bioinformatics web software (Gilchrist and Chooi, 2021). GeneBank format files of putative virulence factors/regulator were uploaded for Bga strain 20GA0385 and *B. glumae* strain BGR1 in the web server to generate the image. The minimum alignment sequence identity threshold was set to 0.3.

### Construction of allelic exchange plasmids

Deletions in strain 20GA0385 were made with pK18*mobsacB* or its Gateway compatible derivative pDONR1k-18ms (AddGene plasmid 72644) (Mijatović et al., 2021). For the unmarked deletion of toxoflavin biosynthesis gene *toxA* and type II secretion apparatus gene *gspC*, 450 bp region upstream and downstream of the respective genes (including start codon with 2 downstream amino acid encoding sequences and stop codon with 2 upstream amino acid encoding sequences) were synthesized as double stranded DNA gblocks with *attB* sites from Twist Biosciences. A unique *Avr*II restriction site was also included in between the flanking region. The synthesized fragment was then BP cloned into pDONR1k18ms plasmid using Gateway BP recombination. The reaction mix was then transformed into chemically competent DH5α cells. The insert in the plasmid was confirmed by restriction digest and sequencing. To generate the T3SS outer membrane protein encoding *sctC* gene deletion mutant, upstream (907 bp) and downstream (790 bp) flanking regions of the ORF was amplified separately using Q5 polymerase PCR reactions. Forward primer amplifying upstream flank and reverse primer amplifying downstream flank were designed with *attB1* and *attB2* sites in the overhangs. Similarly, reverse primer amplifying upstream *sctC* flanking region and forward primer amplifying downstream *sctC* flanking region were designed with overlapping 5’ Gibson overhangs. The Gel extracted purified PCR product was stitched together using Gibson Assembly master mix reaction. The Gibson reaction mix with an expected amplicon size of 1759 bp was excised from the gel, purified, and cloned into pDONR1k18ms using Gateway BP clonase II enzyme mix following manufacturer’s recommendations. The quorum sensing *tofIMR* genes were deleted following a similar strategy as described for the *sctC* gene deletion. The upstream region of *tofI* ORF and downstream region of *tofR* was manually amplified using respective primer pairs tofIMRflankAF2/tofIMRflankAR1 and tofIMRflankBF1/tofIMRflankBR2 with Q5 polymerase-based PCR reaction following manufacturer’s recommendations. Primers tofIMRflankAR1 and tofIMRflankBF1 were designed with 27 bp overlapping sequence region as 5’ overhangs. The overlapping sequence region also included *Kpn*I, *Avr*II, and *Xho*I restriction sites. The two purified PCR products were ligated using overlap extension PCR following conditions and reaction mix as described in Stice et al., 2020. The amplicon size corresponding to the 1636 bp obtained in the gel following overlap extension PCR was excised, purified and cloned into pDONR1k18ms plasmid using Gateway BP Clonase II enzyme reaction. The FRTGm^R^*gus* marker was amplified from the plasmid pCPP5212 was added to the *Xho*I restriction site in between the deletion flanks to aid the selection of single and double crossovers in the downstream steps. The FRTGm^R^*gus* cassette PCR product was excised from the gel, purified, and cloned into *Xho*I digested pDONR1k18ms Δ*tofIMR* deletion construct using NEB T4 ligase reaction following procedures described by Kvitko and Collmer 2012. The ligation reaction mix was transformed into DH5α and the correct insert was confirmed using restriction digest and sanger sequencing. The putative icosalide mutant was created by introducing a premature triple stop codons (TAA) 27 bp downstream of the start codon in the icosalide ORF. The triple stop codon along with a unique *Avr*II restriction site was synthesized along with 450 bp upstream and downstream region in pk18*mobsacB* vector from Twist Biosciences. An independent icosalide interruption mutant was also made mimicking the mutant made by Dose et al., 2018 with some modifications. The authors inserted a kanamycin resistance cassette at the icosalide ORF 7835 bp downstream of the start codon. The upstream (450 bp) and downstream (450 bp) sequence region corresponding to the kanamycin insertion site in Bga strain 20GA0385 along with a unique *Avr*II restriction site in between was synthesized in pk18*mobsacB* plasmid. Instead of kanamycin resistance cassette, the FRTGm^R^*gus* region from the plasmid pCPP5212 was amplified using primer pair icogmgusF and icogmgusR, excised from the gel, and ligated into *Avr*II restriction site using Gibson assembly master mix. For the icosalide and *tofIMR* single crossovers selection, Xgluc and Gm was amended to the LBRfKan medium. The double crossovers were selected following *sacB*-mediated sucrose counter selection. The kanamycin sensitive clones following sucrose counter selection were screened using patch plating. The *toxA* and *gspC* deletion mutants were genotyped using toxAgenoF/toxAgenoR and gspCgenoF/gspCgenoR primers respectively. The *sctC* deletion mutant was confirmed using primer pairs SctCgenoF2/SctCflankBR2 and SctCflankA-F/sctCgenoR. Primer pair SctCgenoF2 and SctCgenoR were designed outside of *sctC* deletion flanks. Similarly, *tofIMR* mutant was confirmed using primer pairs tofIMRgeno-F/GmGUSF and GmGUS-R/tofIMRgeno-R. The primer pair tofIMRgeno-F and tofIMRgeno-R were designed outside of tofIMR deletion flanks. The icosalide FRTGm^R^*gus* interruption mutant was confirmed using primer pairs ico2genoF/GmGUSR2 and GmGUSF/ico2genoR. The ico2genoF and ico2genoR primer pair was designed outside of icosalide deletion flanks. The icosalide triple stop codon interruption mutant was confirmed using primer pair ico1genoF and ico1genoR followed by *Avr*II restriction digest using PCR product as DNA template. Each mutant was also confirmed by partial sangar sequencing of purified PCR product. The *gspC* deletion construct in RHO5 strain was introduced into *sctC* mutant background by biparental mating to generate *gspC sctC* double mutant. Similarly, *toxA* deletion construct was introduced into *gspCsctC* double mutant background by allelic exchange to generate *toxAgspCsctC* triple mutant. The *toxA* mutant was introduced into TTG background to generate *toxATTG* double mutant. Likewise, *gspC* single mutant was introduced into icosalide Gm^R^*gus* mutant background to generate Δ*gspC* Δ*icoS* :: Gm^R^*gus* double mutant.

### Plasmid based complementation vectors

The plasmid-based complementation of single mutants *gspC* and *tofIMR* was done using a broad-host-range expression vector pBBR1MCS-2 (GenBank ID: U02374) and its BP cloning compatible derivative pBBR1MCS2GW (Kovach et al., 1994). The gateway compatible derivative was constructed by first amplifying the gateway cassette from plasmid pR6KT2G using primer pair pBBR1GW-F and pBBR1GW-R. The gel excised and purified PCR amplicon was ligated into *Xba*I digested linearized plasmid using Gibson Assembly master mix reaction. The desalted Gibson reaction was transformed into electrocompetent *ccdB* resistant DB3.1 cells and the transformants were selected on LBKmCam plates. The insert was confirmed using restriction digest and whole plasmid sequencing using PlasmidSaurus. To generate the *gspC* complementation construct, the *gspC* ORF along with 148 bp region was amplified using primer pair gspCcompnF and gspcompnR. The primer pair was designed with *attB* sites overhangs to facilitate Gateway recombination. The PCR gel extracted and purified product was cloned into pBBR1MCS2GW plasmid using Gateway BP clonase reaction. The correct insert was confirmed by genotyping using M13F43 and M13R49 primer pairs. The RHO5 strain with pBBR1MCS2GW plasmid harboring correct *gspC* construct was then conjugated into *gspC* mutant strain using biparental mating. The conjugants selected on LBRfKm plates were confirmed using M13F43 and M13R49 primer-based colony PCR. The PCR amplicon with correct size was excised from the gel, purified and sent for sequencing with M13F primer to confirm the *gspC* insert in the plasmid. The *tofIMR* complementation clone was constructed by PCR amplifying 2225 bp long *tofI*-*tofR* region using primer pair imrcomplnF and imrcomplnR. The purified *tofIMR* PCR product was then cloned into pBBR1MCS2GW plasmid using Gateway BP clonase II reaction. The *tofIMR* insert in the plasmid was confirmed using restriction digest and partial sanger sequencing using M13F primer. The pBBR1MCS2GW plasmid with *tofIMR* complementation construct was transformed to RHO5 cells and mated with *tofIMR* mutant using biparental mating. The conjugants were selected on LBRfKm plates and the correct insert was confirmed by genotyping using M13(−21)F and M13R primer pair.

### Chromosomal restoration of mutants

A chromosomal restoration strategy was then utilized to validate the role of *toxA* gene in yellow pigmentation. A 1691 bp region including the *toxA* ORF and upstream/downstream region was PCR amplified using toxAcomplnF2 and toxAcomplnR2 primer pair containing *attB* sites in the 5’ overhang. The purified PCR product was BP cloned into plasmid pDONR1k18ms and the insert was confirmed using *BsrG*I restriction digest. The RHO5 strain containing the *toxA* restoration construct was then conjugated with *toxA* mutant strain using biparental mating. The sucrose counterselection and patch plating were followed as described above. The *toxA* restoration clone was genotyped by visualizing the yellow pigmentation of the Kan sensitive clones and by colony PCR using primer pair toxAgenocomplnF and toxAgenocomplnR designed outside of the restoration flanks. The *toxA* mutant strain was used as a positive control. The restored clone yielded a bigger amplicon size (1752 bp) in the agarose gel compared to the *toxA* mutant strain (1032 bp). The clone was also confirmed using Sanger sequencing. The *sctC* restoration clone for the *sctC* deletion mutant was obtained using a similar strategy. The *sctC* ORF including 454 bp upstream region and 390 bp downstream region was amplified using primer pair sctCresF and sctCresR containing *attB* sites. The expected amplicon size of 2638 bp was excised from the gel, purified, and cloned into pDONR1k18ms plasmid using BP clonase enzyme mix. The *sctC* insert in the plasmid was confirmed by genotyping using primer pair sctCflankBR and M13R49 and by *BsrG*I restriction digest. The sequence confirmed insert was transformed to RHO5 and conjugated with *sctC* mutant background using biparental mating. Further downstream steps to generate Km sensitive clones were followed as described previously. The kanamycin sensitive clones were screened for mutants using primer pair sctCseqF and sctComplnR using Q5 Polymerase based PCR. The restored clone gave the product size of 2410 bp whereas the *sctC* mutant gave amplicon size of 540 bp.

### Preparation of bacterial inoculum

Bacterial inocula for the casein hydrolysis protease plate assay, *Chromobacterium* AHL assay, tobacco cell death assay, and *in planta* assays were prepared following similar procedure described by Paudel et al., 2024. Briefly, the test strains were streaked on LB or LM plates amended with appropriate antibiotics and incubated at 30^0^C for 24 hours. After 24 hours, the plates were spread with 300 µl of sterile 0.25 mM MgCl_2_/sterile milliQ H_2_0 and incubated overnight at 30^0^C to produce even bacterial lawns. A day after, colony growth from the lawn was scraped and suspended in 1 ml of ddH_2_0 /MgCl_2_ and standardized to OD_600_ = 0.68 - 0.7. The bacterial suspension was serially diluted and plated on LB or LM Rf agar to ensure starting CFU/ml concentration was similar for all the inoculated strains. As the *tofIMR* mutant was gradually reduced in CFU/ml plate recovery, the OD_600_ values for *tofIMR* mutant was variable across the repeats. The standardized suspension was used for inoculation.

### Toxoflavin pigmentation visualization assay

For the toxoflavin pigmentation visualization assay, a 15 ml LB agar media plate was prepared, and sterile nitrocellulose square membranes were placed on top. The membranes were then spotted with 50 µl of overnight cultures and left to dry in the biosafety cabinet. The plate with the dried bacterial suspension was incubated at 30°C for 24 hours. The yellow toxoflavin pigment diffused through the nitrocellulose membrane was visualized on the underside of the plate. The same procedure was followed for visualization of yellow pigment on LM and MGY media as well. For MGY media, 20 ml media volume was used. For a couple of experimental repeats on LM media where the pigmentation was difficult to differentiate between the mutant and WT strain, after taking images after 24 hours, plates were incubated further for 36 hours at room temperature and images were retaken.

### Casein hydrolysis protease plate assay

The casein hydrolysis protease assay was performed for the type II secretion mutant *gspC*, its double and triple mutant derivatives, and the regulatory mutant *tofIMR* following the protocol described by (Brumfield et al., 2018). For the preparation of milk agar plates, 8g of instant skimmed dry milk powder was mixed with sterile milliQ water and autoclaved at 121^0^C for 60 minutes. The second component was prepared by weighing 4.58 g of brain-heart infusion and 6 g agar in 200 ml of sterile milliQ water and autoclaved separately. The milk agar plate was prepared by mixing 15 ml each of the two autoclaved components. The agar was punched by using the back end of a sterile 20 µl pipette tip. The hole was inoculated with 50 µl of normalized bacterial suspension standardized to OD_600_ value of 0.7. The plate was incubated at 30^0^C for 24 hours. Clearing zone on the edge of the inoculated hole indicated the strains were able to hydrolyze casein in the milk powder. The experiment was repeated at least three times.

### Tobacco cell death assay

The tobacco cell death assay was performed with 6–8-week-old tobacco plants (cv. Xanthi SX) grown 26^0^C 12-h day, 22^0^C 12-h night in a Coviron Adaptis growth chamber. At least 3 leaf panels of the third-to eight-oldest leaves were infiltrated with a normalized bacterial suspension of OD_600_ of 0.7 using a blunt syringe. Negative controls were inoculated with 0.25 mM MgCl_2_. The cell death was allowed to develop for at least 3 days before taking the images. Cell death area of the mutants was compared visually to the WT strain. The experiment was repeated at least three times.

### Chromobacterium AHL assay

The AHL reporter *Chromobacterium violaceum* strain CV026 was streaked on a LM agar plate to make a lawn and incubated at 30^0^C for 48 hours. A dense suspension of CV026 was prepared by scraping the bacterium from the lawn into sterile milliQ H_2_0. A bacterial lawn of CV026 was prepared on LM agar plate by using 300 µl of dense CV026 suspension. After letting it dry, the agar was punched with back end of a sterile 20 µl pipette tip to make holes. Normalized Bga suspension of 50 µl with OD_600_ = 0.7 was inoculated into the holes and incubated at 30^0^C for 48 hours. The inoculated holes were monitored for the appearance of violet ring around the edges produced by the reporter strain. The CV026 bacterial suspension was used as negative control. The experiment was repeated three times on its entirety.

### Swarming motility assay

Swarming motility plate assay was performed following the protocol as described by (Dose et al., 2018) with slight modifications. Briefly, MGY media was prepared by mixing 1.5 g glycerol, 0.188 g yeast extract and 1.5% agar in 144 ml milliQ water and autoclaved at 121^0^C for 60 minutes. M9A salt was prepared by mixing 2.50 g K_2_HPO_4_ and 1 g KH_2_PO_4_ in 100 ml of milliQ water and sterilized by autoclaving. Similarly, M9B was prepared by mixing 2.58 g of sodium citrate dihydrate, 5 g of (NH_4_)_2_SO_4_ and 0.5 g of MgSO_4._ 7H_2_O in 100 ml of milliQ water. MGY agar media was prepared the same day of inoculation. After autoclaving, 6 ml each of warm M9A and M9B salt was mixed to MGY agar media and diluted in half with MGY broth and a 20 ml mix was poured into the plate. To prepare the bacterial suspension, WT and mutants were streaked to isolation on LB Rf or LB Rf Kan plates and a single colony was used to start overnight cultures (∼24 h). A normalized bacterial suspension of 0.7 OD_600_ was prepared using sterile 0.25 mM MgCl_2_ and 50 µl was inoculated into the center of the plate. Suspension was allowed to dry and incubated at 28^0^C for 21 hours. At least one plate was plated for each strain and the experiment was repeated three times in its entirety.

### Onion foliar assay

Onion plants (*Allium cepa* L. cv Century) 1.5 to 4.5 months old grown under greenhouse conditions from February to May were used for inoculation. The procedure for inoculation, sampling and bacterial quantification was followed as described by (Paudel et al., 2024b). Inoculum for Bga WT strain 20GA0385 and its mutant derivatives were prepared as described above. Onion plants were trimmed to keep the two oldest and tallest leaves intact. Two leaves per plant and three biological replicates per strain were inoculated. The approximate midpoint and 0.5 cm length top and bottom was marked for processing and the midpoint was poked/wounded with a sterile pipette tip on one side to drop the inoculum. Each leaf was inoculated with 10 µl of a normalized bacterial suspension. Two leaves were inoculated with 0.25 mM MgCl_2_ as negative controls. The maximum necrosis length was measured 3 days post inoculation. The experiment was repeated at least three times in its entirety.

For quantification of bacterial population in the infected blade tissue, a pre-marked 0.5 cm section above and below the inoculation site was excised with a sterile scissor and added to 200 µl of Milli-Q H_2_0 in a 2-ml SARSTEDT microtube (SARSTEDT AG & Co., Numbrecht, Germany) along with three 3-mm zirconia beads (Glen Mills grinding media). The tissue was crushed using a Bead Ruptor Elite Bead Mill Homogenizer (Omni International, Kennesaw, GA) for 30 seconds at 4 m/s speed settings. Serial dilutions were performed by mixing 20 µl of the ground tissue with 180 µl of sterile 0.25 mM MgCl_2_ in 96-well styrene plates. Diluents were plated on LB or LM plates amended with rifampicin. The number of colony-forming units (CFUs) was back-calculated to determine the bacterial population per mg of the infected tissue. The recovered CFU/mg values for two leaves per plant was averaged making it a total of three data points for three biological replicates per treatment.

### Onion seedling assay

Onion seedling assay was done to determine the contribution of Δ*toxA*Δ*gspC*Δ*sctC* triple mutant to seedling necrosis and bacterial populations. Onion seedlings grown for 6-8 months were used for inoculation. One leaf per plant per pot was selected for inoculation. A normalized bacterial suspension of 0.7 OD_600_, 10 µl was used to inoculate the mid-point of the marked seedlings. Bacterial CFU/ml plate recovery of the WT and mutant strains were determined on day 0 to ensure starting concentration was similar. The seedling necrosis length and in planta population counts were assessed at 3 dpi, following the protocol detailed in the onion foliar assay section above.

### Red Scale Necrosis (RSN) assay

The RSN assay was done following the procedure described in Stice et al., 2018; Shin et al., 2023; Paudel et al., 2024b with slight modifications. The store brought red onion bulbs were cut into 3-5 cm x 3-5 cm sized scales. The scales were sterilized in a 3 % Sodium hypochlorite/Clorox solution and washed several times with deionized H_2_0. Six scales per strain were kept on top of a sterile pipette tip rack in a flat floored with wetted paper towels. The scales were wounded on the center with a pipette tip. The wounded scale was deposited with 10 µl of normalized bacterial suspension (OD_600_ = 0.82 for *tofIMR* and 0.68 – 0.7 for all other treatments). To ensure all the tested strains had comparable CFU/ml plate recoveries, 2 to 3 replicates of normalized bacterial suspension were plated on LB or LM media amended with rifampicin following serial dilutions. Strains that had comparable CFU/ml reading were further utilized for the experiment. The scales were incubated at room temperature for 3 days. Necrosis area around the point of inoculation was measured using ImageJ software. The experiment was repeated at least three times for each strain. For the quantification of bacterial populations from the inoculated scales, onion tissues were sampled at 3 dpi at the point of inoculation using ethanol sterilized metal borer (*r =* 2.5 mm). The sample was processed for crushing and serial dilutions to calculate the CFU/mg for each scale following the procedure as described in the above section.

For mutants that exhibited significantly smaller area of red scale necrosis area compared to the WT strain, additional rounds of RSN assay were performed with a 100-fold diluted bacterial suspension of 0.7 OD_600_ density. CFU/ml recovery of 0.7 OD_600_ was calculated for each treatment to ensure the starting bacterial concentration was comparable for all treatments. The red scale assay procedure for inoculation, sampling, and *in planta* bacterial quantification was followed as described before.

### Data analysis and presentation

The statistical difference in scale/foliar/seedling necrosis area and bacterial populations in Log (CFU/mg) between the WT and mutants was performed using pairwise t-test function applying the Bonferroni coefficient in R studio 2023.09.0. For treatments that had an unequal number of observations, Welch’s t-test was performed with t.test function in R studio 2023.09.0 using “var.equal = FALSE” option.

The box plot was generated using ggplot function in R studio. The color of the box was assigned based on function. The WT strain was represented in green, secretion system mutants in grey, and mutants for the metabolites toxoflavin and icosalide in yellow, while regulator mutant *tofIMR* was assigned red. Double and triple mutants were given distinct colors: the *toxA* TTG mutant was marked in white, Δ*gspC*Δ*sctC* in cyan, Δ*gspC*Δ*icoS*::Gm^R^*gus* in orange, and the triple mutant *toxA*/*gspC*/*sctC* was represented in purple.

## Acknowledgements

We acknowledge all the members of Kvitko Lab, Dutta Lab, and Yang lab for their helpful comments and suggestion during the preparation of this manuscript.

---

**Supplementary Table S1:** Amino acid identity % between the putative virulence factors/regulators in *B. glumae* strain BGR1 and Bga strain 20GA0385. Data generated using Clinker CAGECAT Web server.

| Putative virulence factors | % aa identity |
| --- | --- |
| Toxoflavin (biosynthesis cluster) (*B. glumae* BGR1 vs Bga 20GA0385) |  |
| ToxA | 96 |
| ToxB | 98 |
| ToxC | 97 |
| ToxD | 96 |
| ToxE | 90 |
| Type II secretion system (*B. glumae* BGR1 vs Bga 20GA0385) |  |
| GspD | 86 |
| GspE | 94 |
| GspF | 96 |
| GspC | 90 |
| GspG | 97 |
| GspH | 88 |
| GspI | 81 |
| GspJ | 76 |
| GspK | 78 |
| GspL | 84 |
| GspM | 83 |
| GspN | 81 |
| Type III secretion system (*B. glumae* BGR1 vs Bga 20GA0385) |  |
| SctS | 94 |
| SctR | 97 |
| SctQ | 70 |
| HpaP/SctP | 75 |
| SctV | 91 |
| SctU | 90 |
| SctG | 88 |
| HrpB2 | 91 |
| SctJ | 89 |
| HrpB4 | 84 |
| SctL | 71 |
| SctN | 94 |
| HrpB7 | 68 |
| SctT | 84 |
| AraC | 93 |
| SctC | 85 |
| HrpW putative effector | 59 |
| Putative Response regulator | 85 |
| Icosalide (Bga 20GA0385 vs *B. gladioli* HKI0739) |  |
| IcoS | 97 |
| Quorum sensing (*B. glumae* BGR1 vs Bga 20GA0385) |  |
| TofI | 87 |
| TofM | 80 |
| TofR | 92 |

**Supplementary Table S2:**
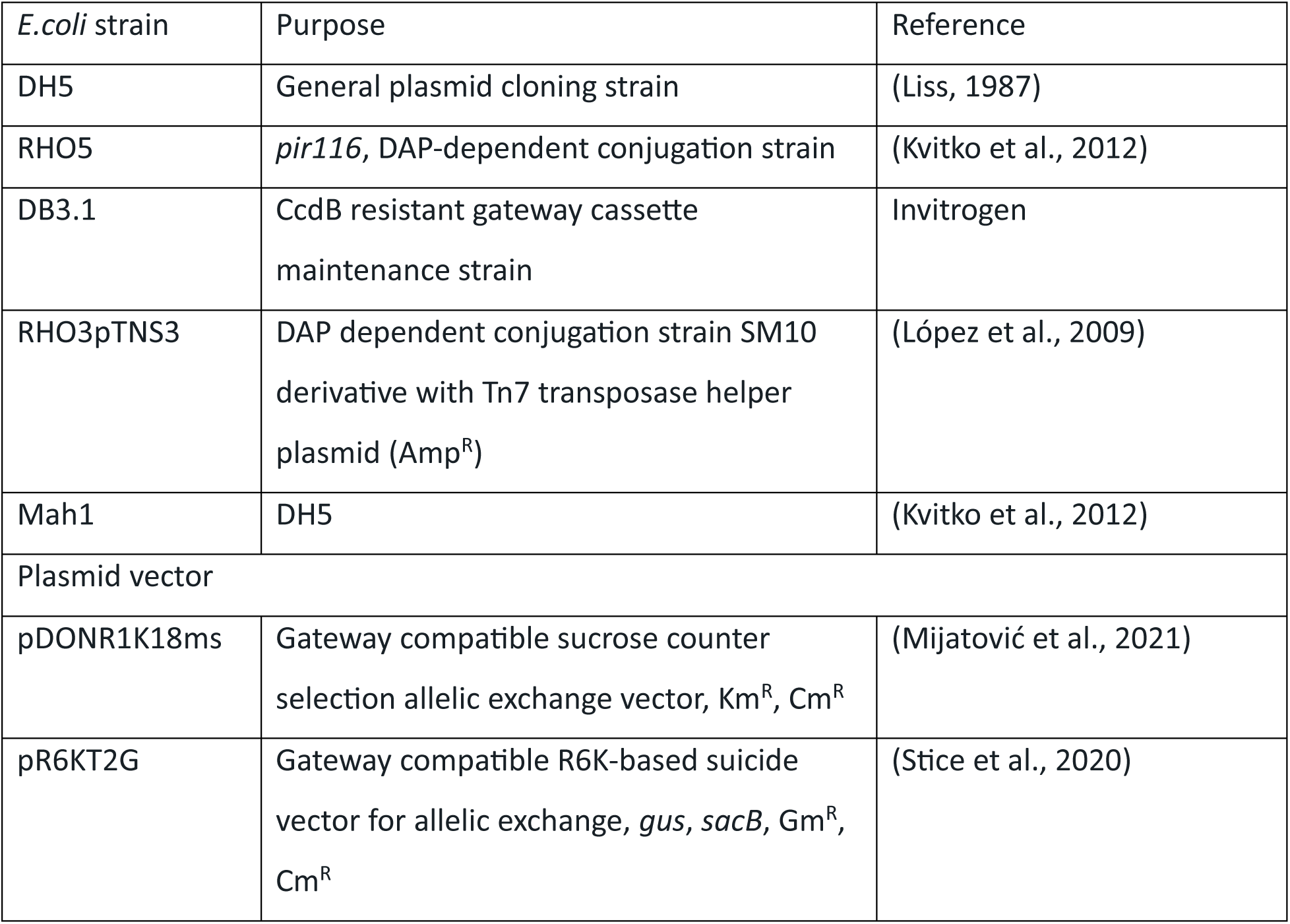
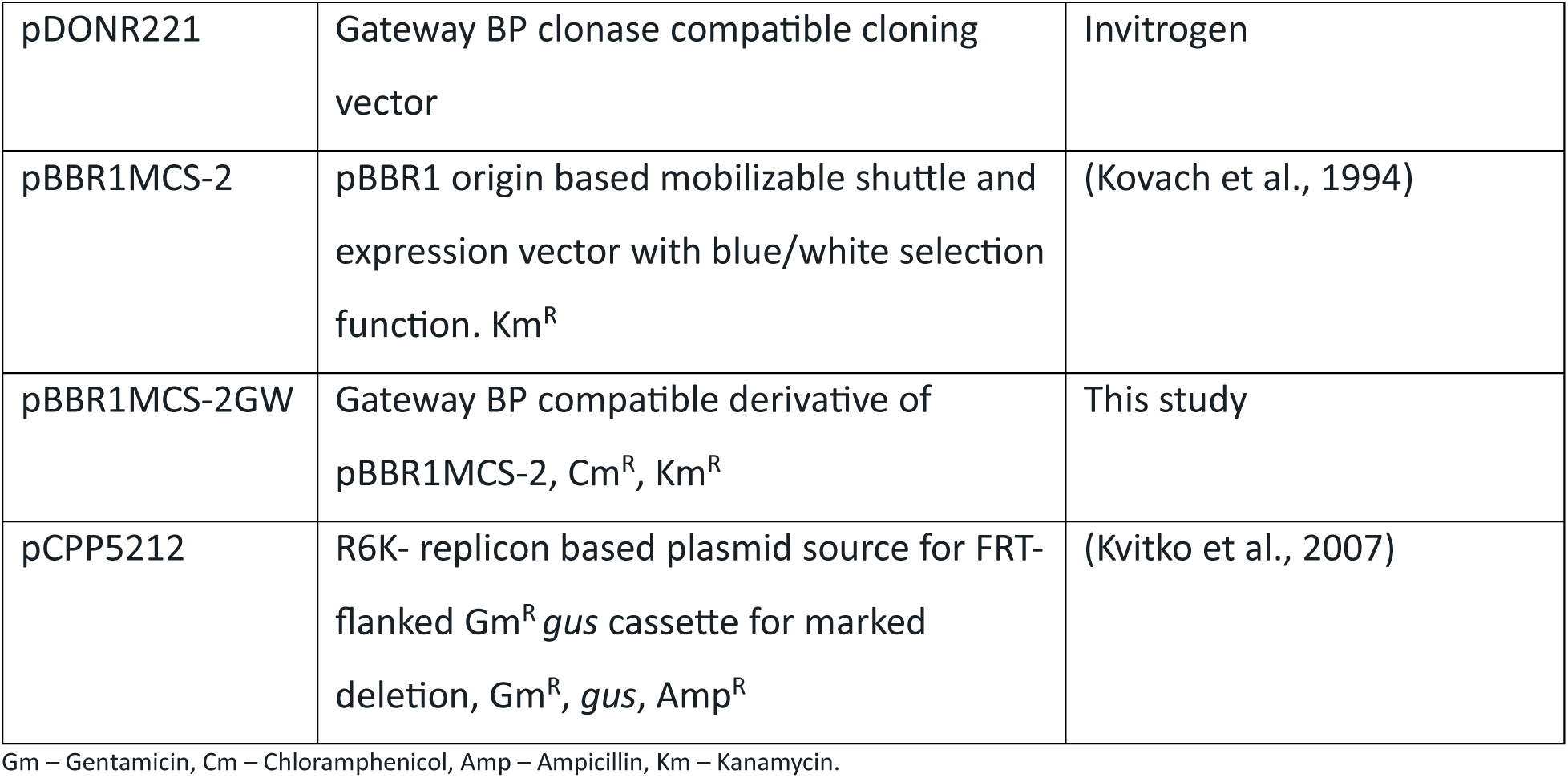
*E. coli* genotype and plasmids used in the study.

| *E. coli* strain | Purpose | Reference |
| --- | --- | --- |
| DH5 | General plasmid cloning strain | (Liss, 1987) |
| RHO5 | *pir116*, DAP-dependent conjugation strain | (Kvitko et al., 2012) |
| DB3.1 | CcdB resistant gateway cassette maintenance strain | Invitrogen |
| RHO3pTNS3 | DAP dependent conjugation strain SM10 derivative with Tn7 transposase helper plasmid (Amp<sup>R</sup>) | (López et al., 2009) |
| Mah1 | DH5 | (Kvitko et al., 2012) |
| Plasmid vector |  |  |
| pDONR1K18ms | Gateway compatible sucrose counter selection allelic exchange vector, Km<sup>R</sup>, Cm<sup>R</sup> | (Mijatović et al., 2021) |
| pR6KT2G | Gateway compatible R6K-based suicide vector for allelic exchange, *gus*, *sacB*, Gm<sup>R</sup>, Cm<sup>R</sup> | (Stice et al., 2020) |
| pDONR221 | Gateway BP clonase compatible cloning vector | Invitrogen |
| pBBR1MCS-2 | pBBR1 origin based mobilizable shuttle and expression vector with blue/white selection function. Km<sup>R</sup> | (Kovach et al., 1994) |
| pBBR1MCS-2GW | Gateway BP compatible derivative of pBBR1MCS-2, Cm<sup>R</sup>, Km<sup>R</sup> | This study |
| pCPP5212 | R6K- replicon based plasmid source for FRT-flanked Gm<sup>R</sup> *gus* cassette for marked deletion, Gm<sup>R</sup>, *gus*, Amp<sup>R</sup> | (Kvitko et al., 2007) |

Gm – Gentamicin, Cm – Chloramphenicol, Amp – Ampicillin, Km – Kanamycin.

**Supplementary Table S3:**
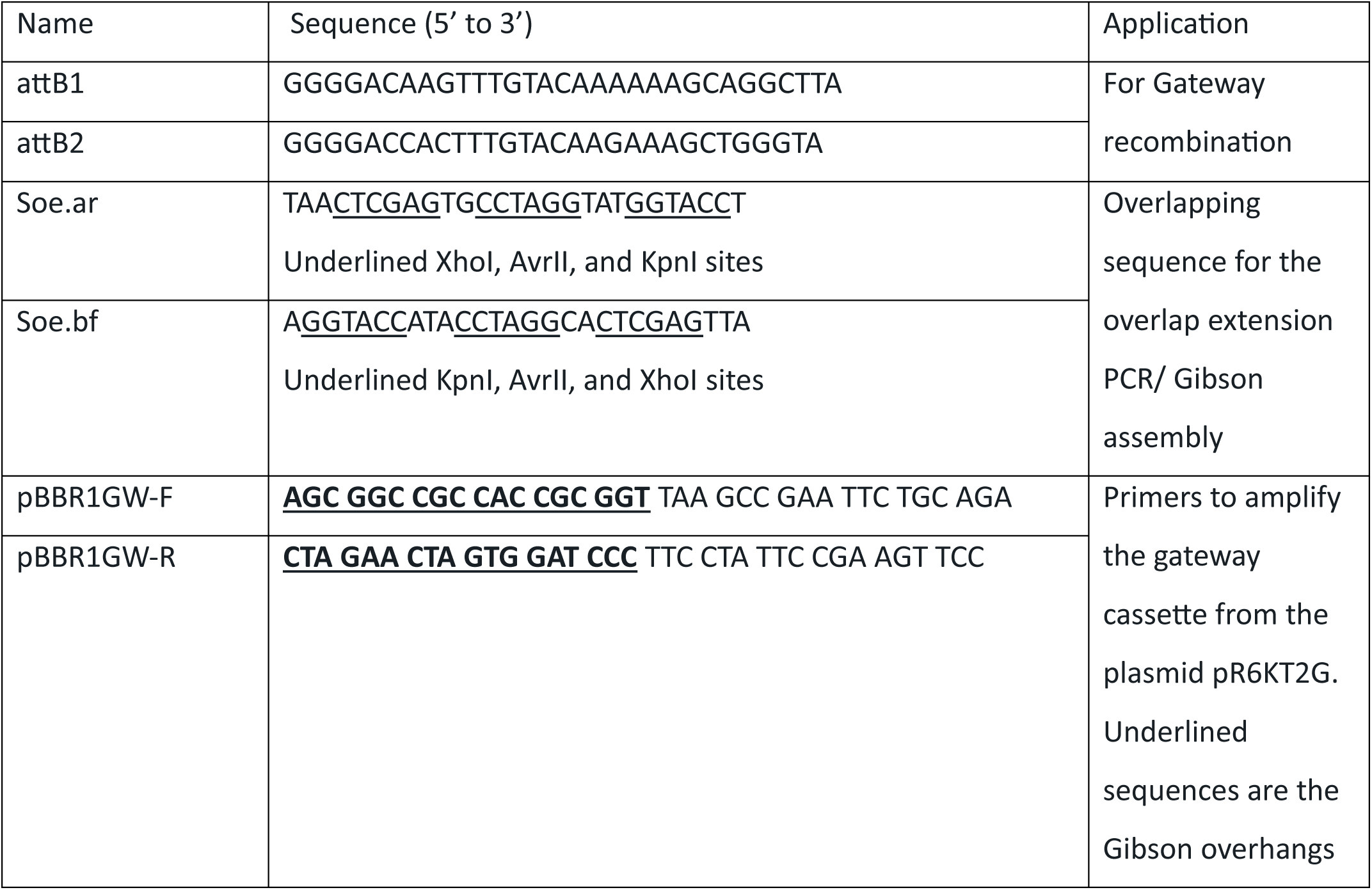
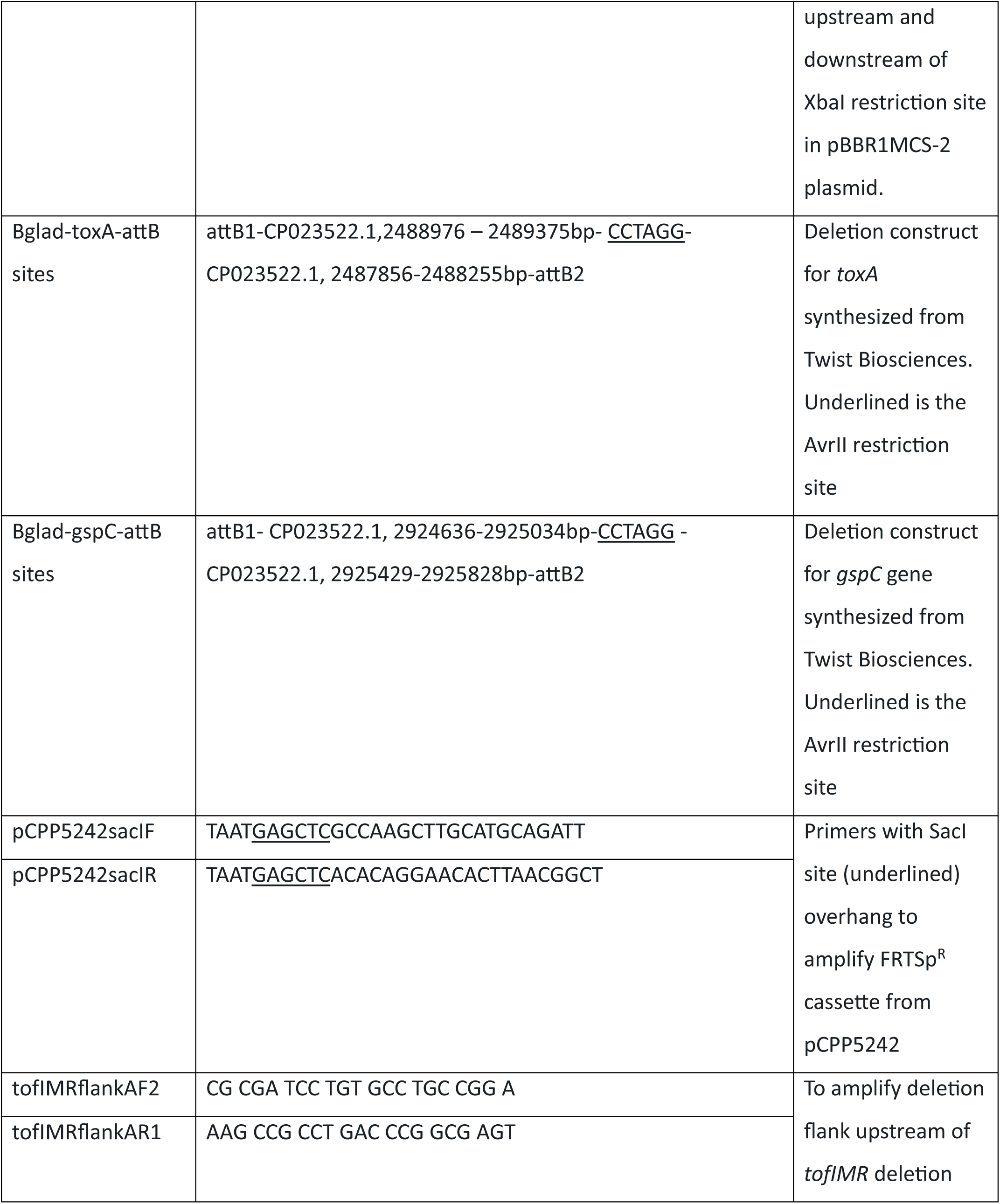
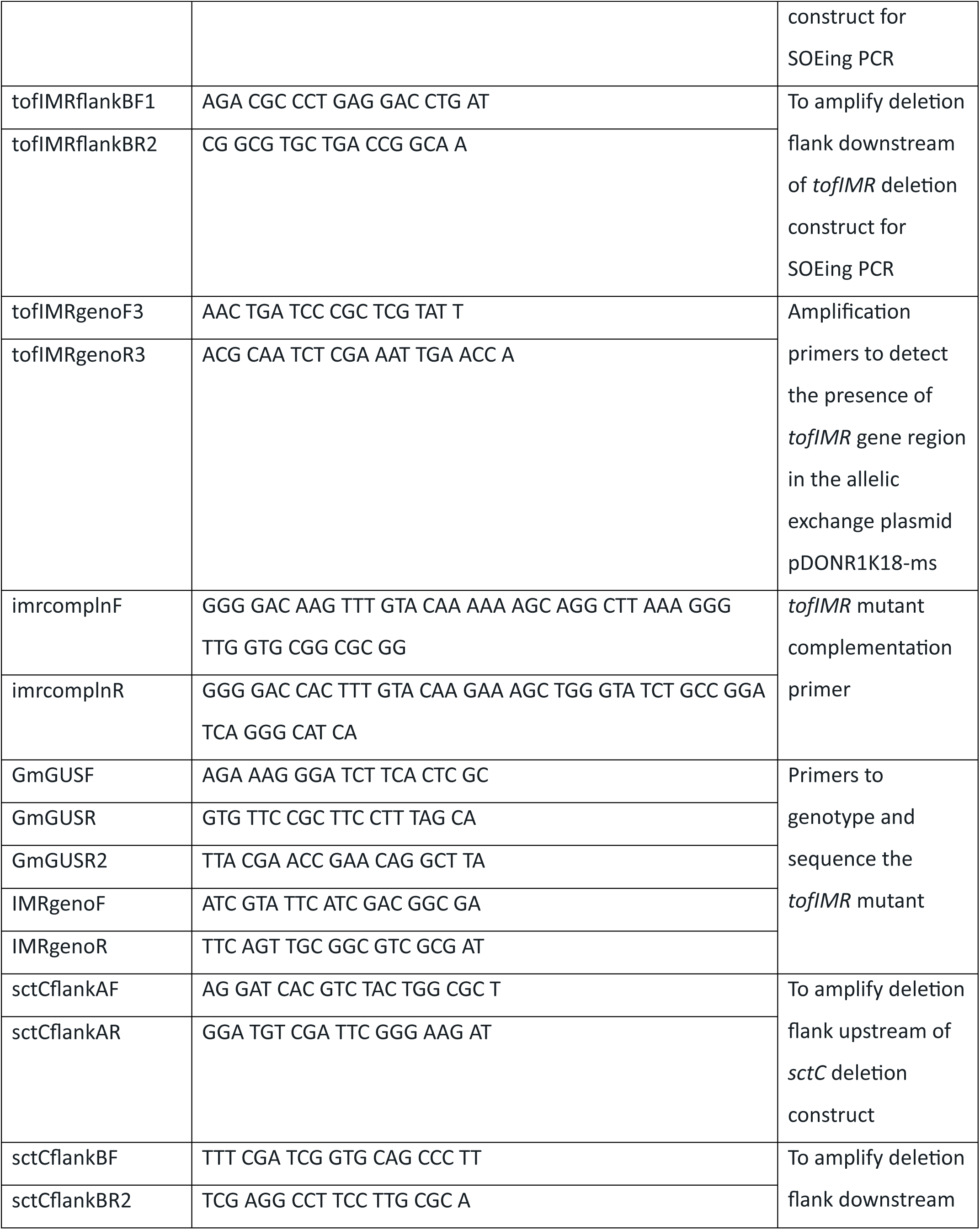
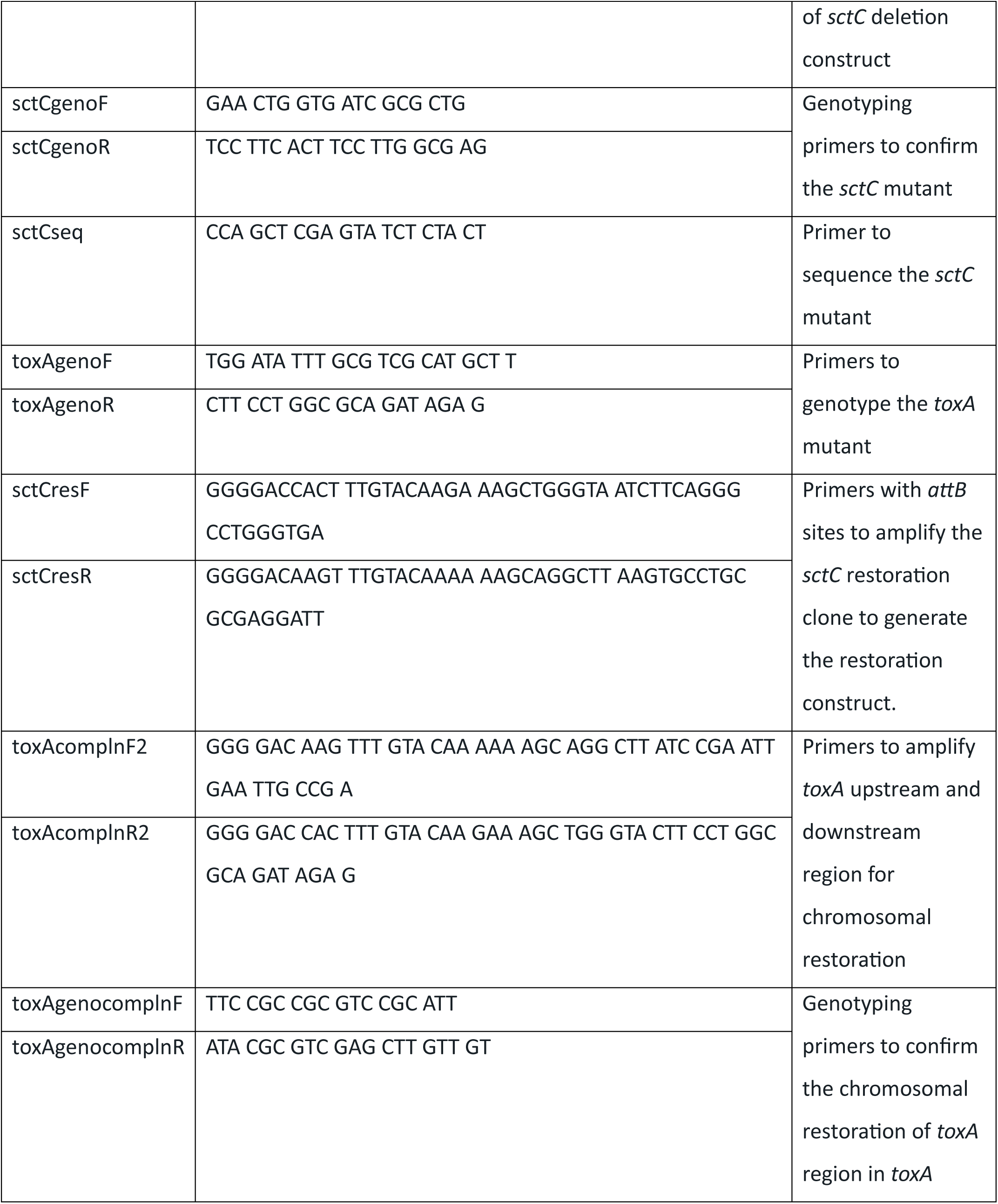
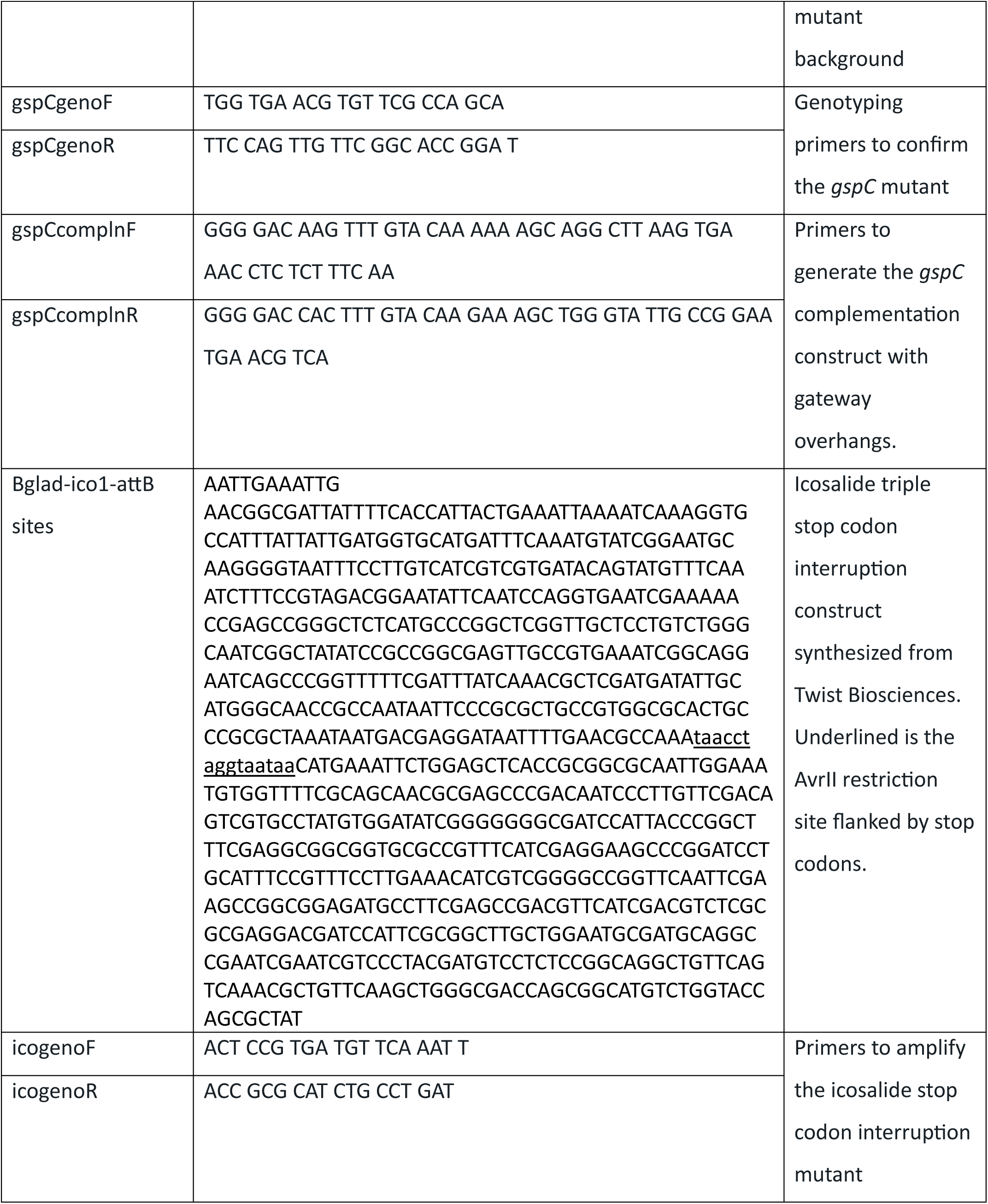
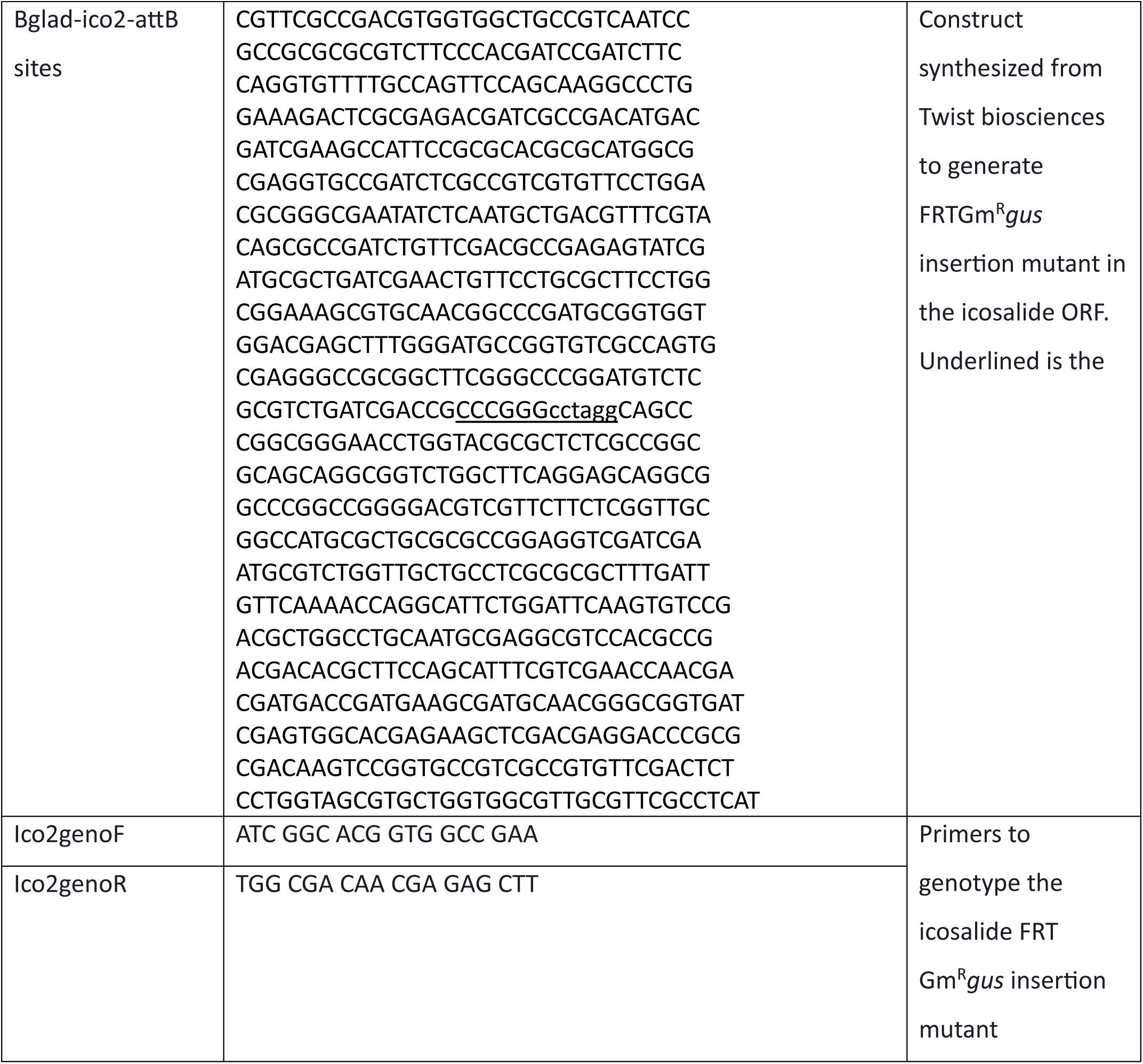
Primers/Gene fragments used in this study for allelic exchange, sequencing, and PCR.

**Supplementary Table S4:**
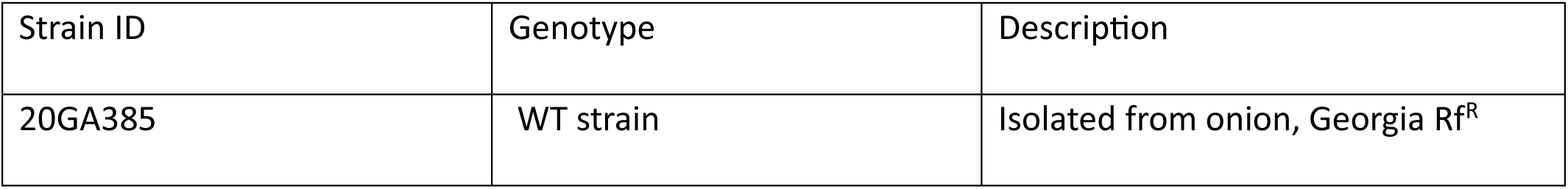
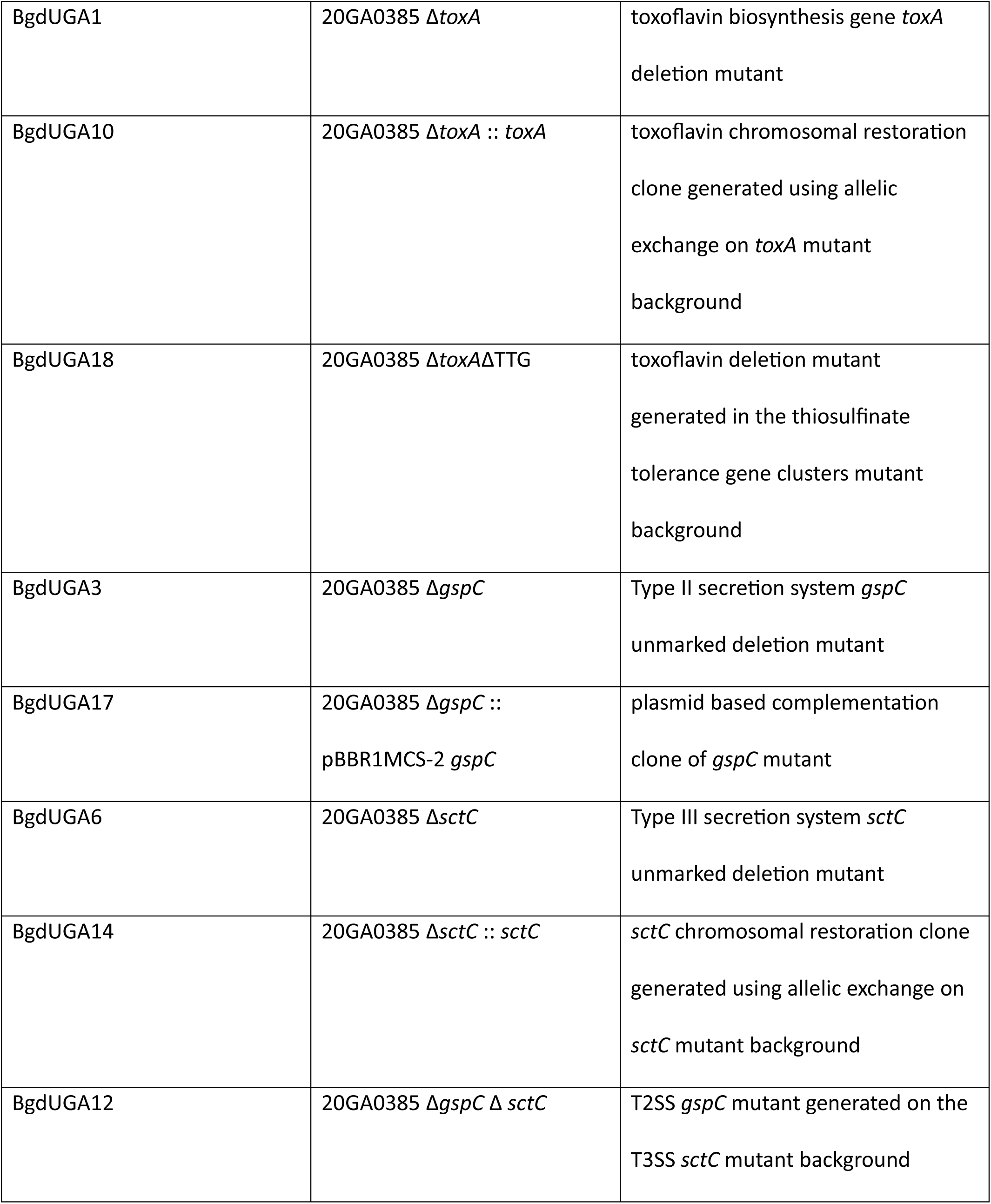
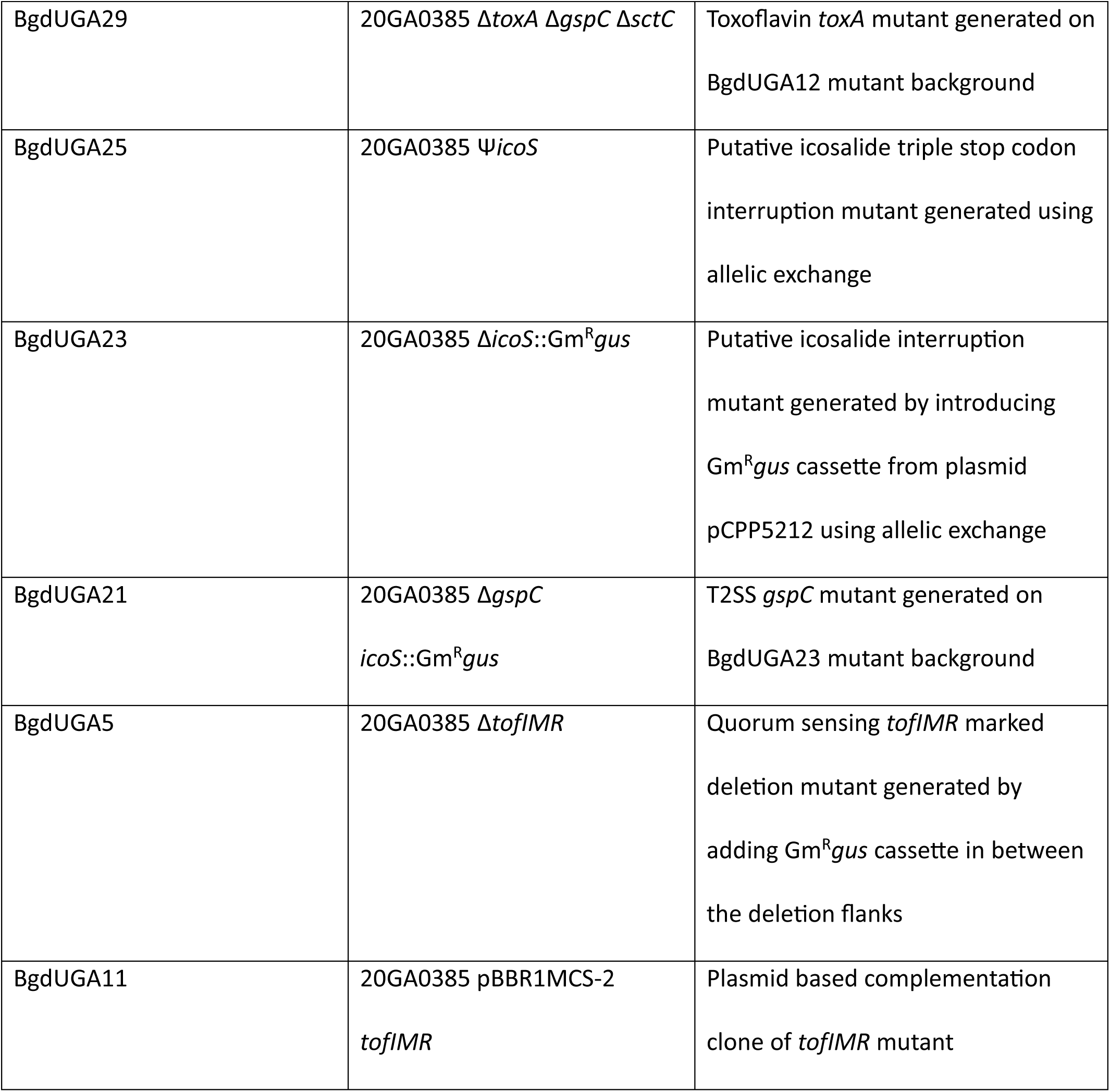
*Burkholderia* strains and mutants generated in the study.

| Strain ID | Genotype | Description |
| --- | --- | --- |
| 20GA385 | WT strain | Isolated from onion, Georgia Rf<sup>R</sup> |
| BgdUGA1 | 20GA0385 Δ*toxA* | toxoflavin biosynthesis gene *toxA*<br>deletion mutant |
| BgdUGA10 | 20GA0385 Δ*toxA* :: *toxA* | toxoflavin chromosomal restoration<br>clone generated using allelic<br>exchange on *toxA* mutant<br>background |
| BgdUGA18 | 20GA0385 Δ*toxA*ΔTTG | toxoflavin deletion mutant<br>generated in the thiosulfinate<br>tolerance gene clusters mutant<br>background |
| BgdUGA3 | 20GA0385 Δ*gspC* | Type II secretion system *gspC*<br>unmarked deletion mutant |
| BgdUGA17 | 20GA0385 Δ*gspC* ::<br>pBBR1MCS-2 *gspC* | plasmid based complementation<br>clone of *gspC* mutant |
| BgdUGA6 | 20GA0385 Δ*sctC* | Type III secretion system *sctC*<br>unmarked deletion mutant |
| BgdUGA14 | 20GA0385 Δ*sctC* :: *sctC* | *sctC* chromosomal restoration clone<br>generated using allelic exchange on<br>*sctC* mutant background |
| BgdUGA12 | 20GA0385 Δ*gspC* $\Delta$ *sctC* | T2SS *gspC* mutant generated on the<br>T3SS *sctC* mutant background |
| BgdUGA29 | 20GA0385 $\Delta toxA \Delta gspC \Delta sctC$ | Toxoflavin *tox*A mutant generated on BgdUGA12 mutant background |
| BgdUGA25 | 20GA0385 $\Psi icoS$ | Putative icosalide triple stop codon interruption mutant generated using allelic exchange |
| BgdUGA23 | 20GA0385 $\Delta icoS::Gm^Rgus$ | Putative icosalide interruption mutant generated by introducing *Gm<sup>R</sup>gus* cassette from plasmid pCPP5212 using allelic exchange |
| BgdUGA21 | 20GA0385 $\Delta gspC$<br>*icoS::Gm<sup>R</sup>gus* | T2SS *gspC* mutant generated on BgdUGA23 mutant background |
| BgdUGA5 | 20GA0385 $\Delta tofIMR$ | Quorum sensing *tofIMR* marked deletion mutant generated by adding *Gm<sup>R</sup>gus* cassette in between the deletion flanks |
| BgdUGA11 | 20GA0385 pBBR1MCS-2<br>*tofIMR* | Plasmid based complementation clone of *tofIMR* mutant |

**Supplementary Figure S1: Putative virulence factors and regulator are conserved in *B. gladioli* strain 20GA0385 and *B. glumae* strain BGR1.** Clinker gene synteny arrow diagram showing the gene organization and synteny of putative virulence factors/regulator - **A,** toxoflavin biosynthesis gene cluster, **B,** quorum sensing *tofIMR* system**, C**, type II secretion system, and **D,** type III secretion system. Genes are color coded based on predicted function. The percentage of amino acid identity between the linked genes is shown in gradient scale. The annotation of genes in the cluster is used from CAGECAT web server platform. Image generated using Clinker CAGECAT web server. Toxoflavin and homo serine lactones structures used from the internet.

**Supplementary Figure S2: Functional validation of select mutants utilized in the study.** Visualization of diffusible yellow toxoflavin pigmentation from the back side of plate in the nitrocellulose membrane spotted on top of **A,** MGY agar plate, **B, C** LM agar plate with (from left to right: WT, quorum sensing mutant *tofIMR*, and *tofIMR* complementation clone. The plate from Exp 2 and Exp 3 was further incubated at room temperature and the image was captured after 2 and half days of inoculation. Representative image from one of the experimental repeats is shown for panel A. **D,** Casein protease plate assay showing the clearing zone phenotype of tested (from left to right) WT, Δ*gspC*Δ*sctC*, Δ*toxA*Δ*gspC*Δ*sctC* and Δ*gspC*Δ*icoS*::Gm^R^*gus*. n = total number of observations. Representative image from experimental repeats is shown. **E**, Tobacco cell death assay phenotype of different mutants compared to the WT strain. Representative image from one of the experimental repeats is shown. The ratio of infiltrated spots displaying the representative image phenotype to the total number of infiltrated spots is shown at the bottom of each strain. **F,** swarming phenotype of the tested mutants compared to the WT strain. Image taken after 21 hours. Representative image from one of the experimental repeats is shown. n is the number of independent experiments. Scale: 1 cm

**Supplementary Figure S3:** Box plot showing the red scale in planta population count of WT strain compared to **A,** Δ*toxA* **B,** Ψ*icoS***, C,** Δ*icoS*::Gm^R^*gus***, D,** Δ*toxA*ΔTTG**, E,** Δ*gspC* **F,** Δ*sctC* **G,** Δ*gspC*Δ*sctC***, H,** Δ*gspC*Δ*icoS*::Gm^R^*gus*, **I**, Δ*toxA*Δ*gspC*Δ*sctC*, **J,** Δ*tofIMR***, K,** Δ*gspC***, L,** Δ*gspC*Δ*sctC***, M,** Δ*gspC*Δ*icoS*::Gm^R^*gus* **N,** Δ*toxA*Δ*gspC*Δ*sctC*, and Δ*tofIMR* mutants. n is the total number of observations. 100-fold label in the **Figures K-O** represents that the treatments were compared at 100-fold diluted starting bacterial concentration. The test of significance between the log fold bacterial populations was performed using pairwise t-test function in RStudio.

**Supplementary Figure S4:** Box plot showing the comparison of **A,** foliar necrosis length and **B,** in planta bacterial populations between the WT and *toxA* restoration clone. Box plot showing the comparison of in planta foliar bacterial populations of WT compared to **C,** Δ*gspC*, **D,** Δ*sctC*, and **E,** Δ*gspC*Δ*sctC*. A representative image of infected leaves or scales inoculated with treatments at 3 dpi is presented above the box plot where applicable. n is the total number of observations across the experimental repeats.

