## Supplementary figures and images for "Distinct Virulence Mechanisms of *Burkholderia gladioli* in Onion Foliar and Bulb Scale Tissues"

### Supplementary Figure S1

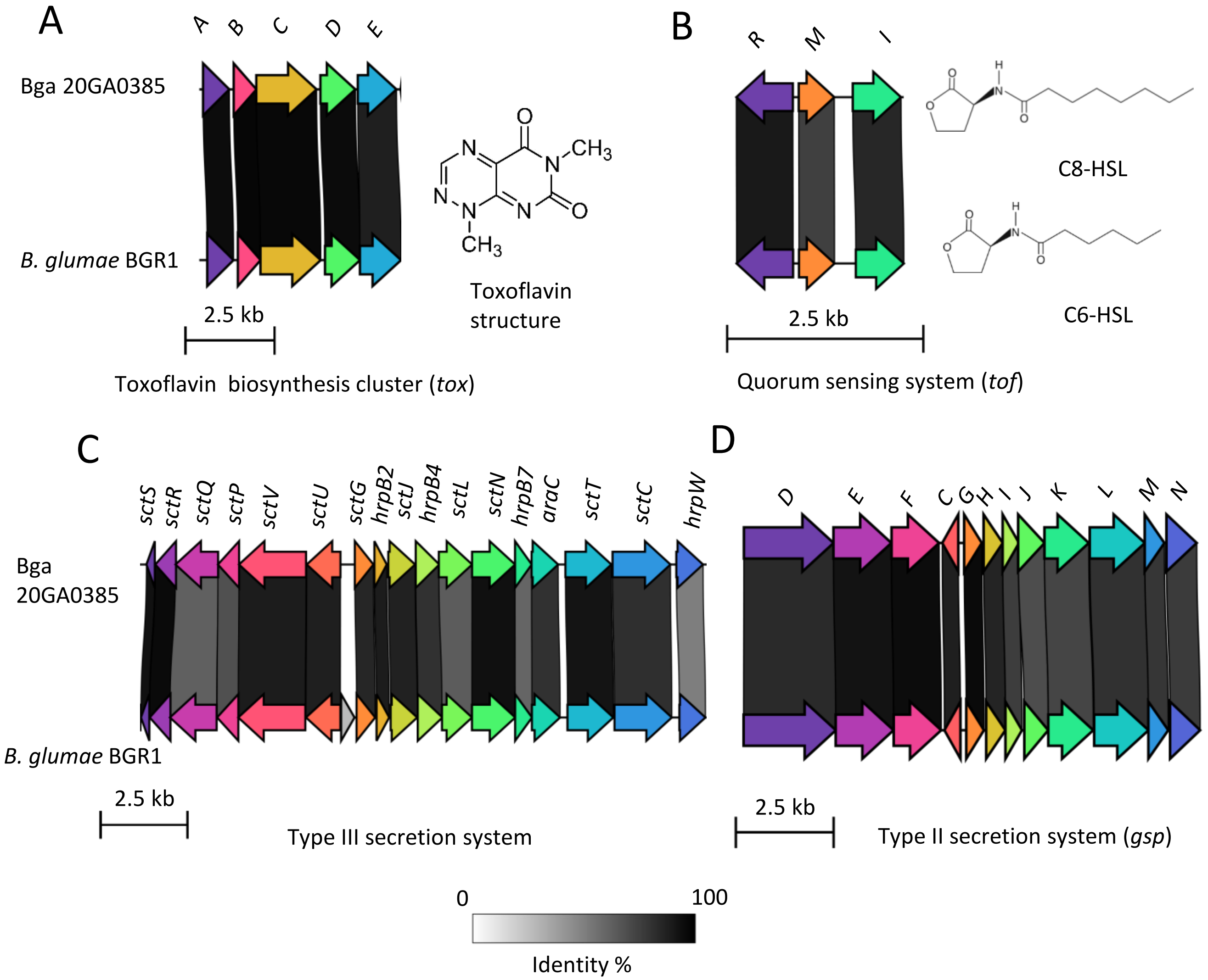

### Supplementary Figure S2

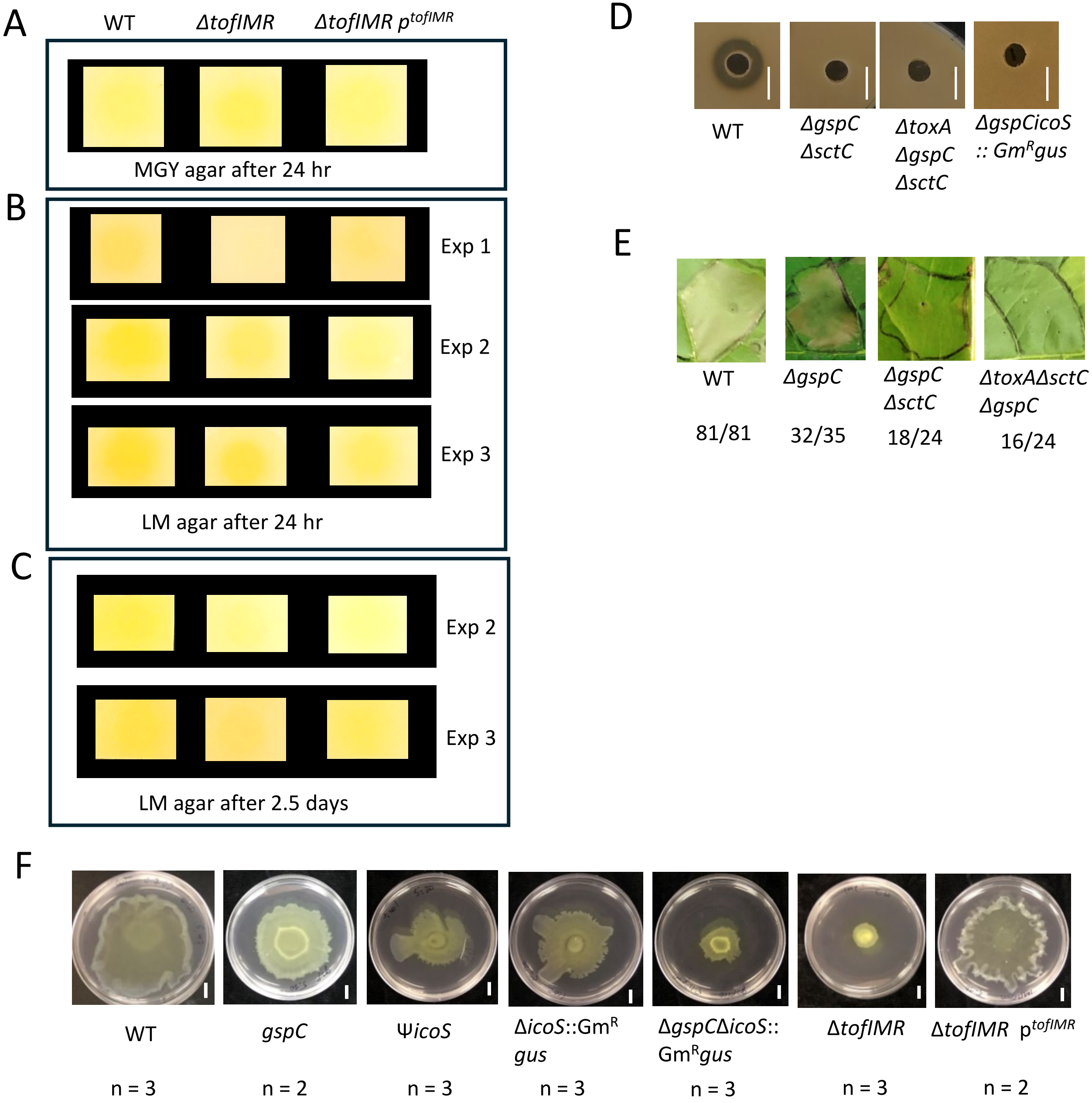

### Supplementary Figure S3

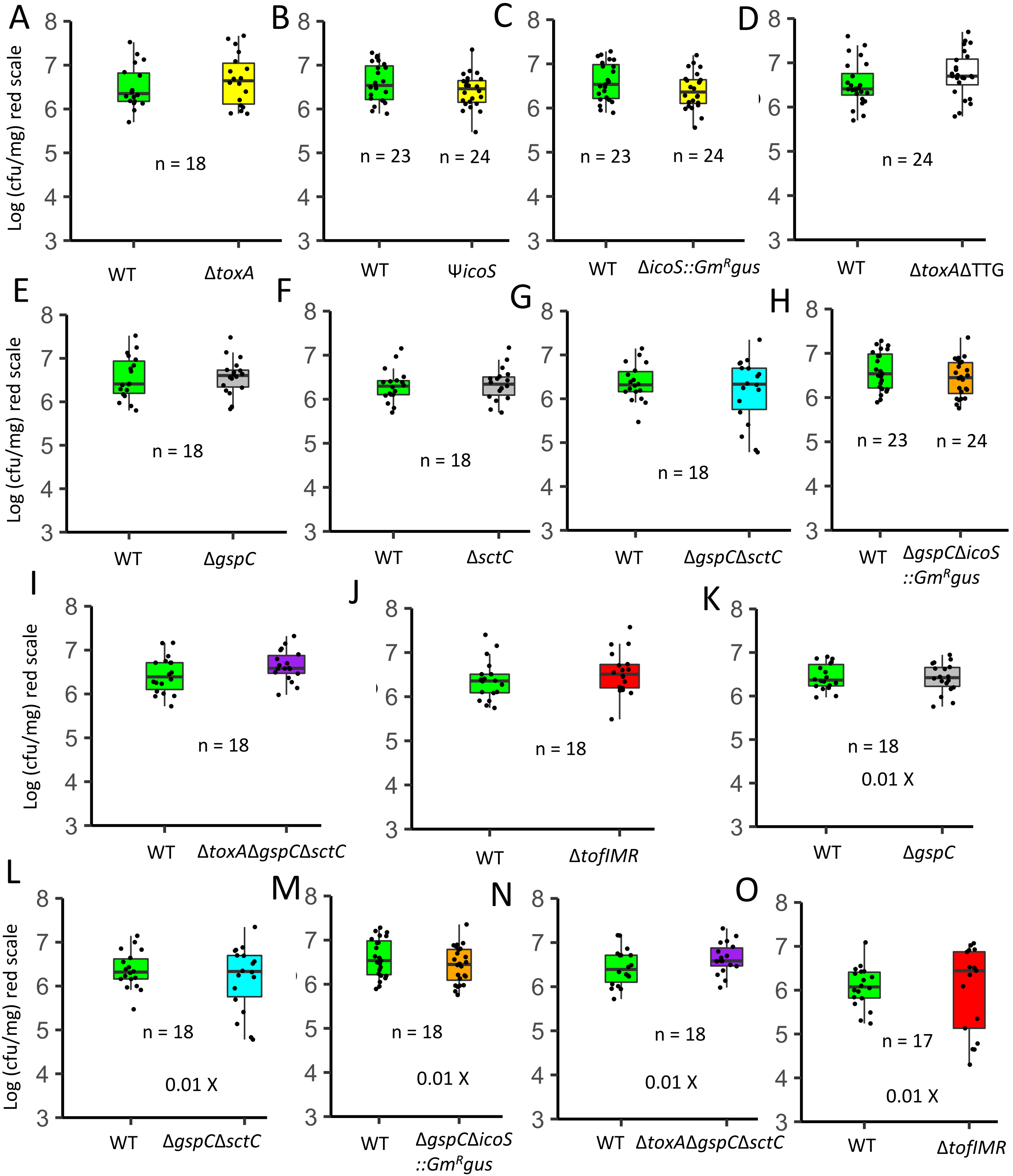

### Supplementary Figure S4

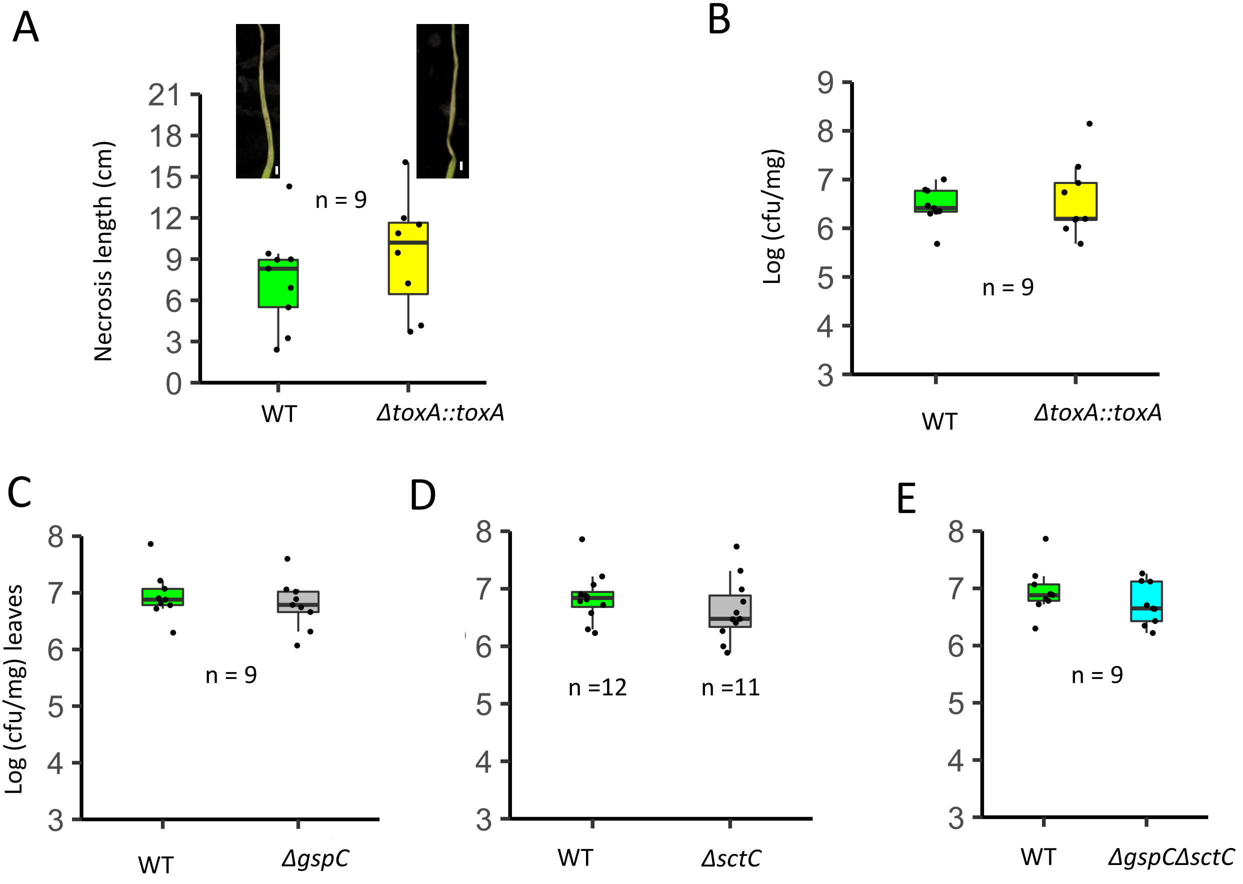
